# Bayesian arrival model for Atlantic salmon smolt counts powered by environmental covariates and expert knowledge

**DOI:** 10.1101/399618

**Authors:** Henni Pulkkinen, Panu Orell, Jaakko Erkinaro, Samu Mäntyniemi

## Abstract

Annual run size and timing of Atlantic salmon smolt migration was estimated using Bayesian model framework and data from six years of a video monitoring survey. The model has a modular structure. It separates sub-processes of departing, traveling and observing, of which the first two together define the arrival distribution. The sub-processes utilize biological background and expert knowledge about the migratory behavior of smolts and about the probability to observe them from the video footage under varying environmental conditions. Daily mean temperature and discharge were used as environmental covariates. The model framework does not require assuming a simple distributional shape for the arrival dynamics and thus also allows for multimodal arrival distributions. Results indicate that 20% - 43% of smolts passed the Utsjoki monitoring site unobserved during the years of study. Predictive studies were made to estimate daily run size in cases with missing counts either at the beginning or in the middle of the run, indicating good predictive performance.

## Introduction

Migratory fish species are often monitored along their migration routes at different life stages to collect fishery-independent data for stock assessment. Monitoring methods include e.g. counting fences, video, sonar and snorkeling counts, and common to all is that they rarely provide perfect information about the number of individuals passing the system (Dempson et al. 1991; Romakkaniemi et al. 2000; Orell et al. 2007; 2011). Part of the fish run may be missed because of difficult environmental conditions or device failures, there may be double counting, or the data achieved may be otherwise partial or biased (e.g. Holmes et al. 2006). Thus, the assumptions made when interpreting the data can have a great influence on the estimated stock abundance.

Increasing the information beyond the actual monitoring data collected can improve the trustworthiness of stock assessments (Kuparinen et al. 2012): such information can be based on the resources available in other fields of biological research such as ecology or life history theory, but also on the physical and technical details of the observation processes. Methods of Bayesian inference enable both estimation of uncertainty and combination of various sources of information. Statistical methods are often used, however, without considering the biological realism and just focusing on statistical data analysis. Such a procedure easily leads to awkward model assumptions that cannot be biologically interpreted and the potential effects of those modelling decisions may pass undiscussed. We echo the idea of Kuparinen et al. (2012) and Mäntyniemi et al. (2015) in asserting that biological plausibility has the priority over measures of statistical fit and information theoretic parsimony when it comes to assessing the performance of stock assessment models.

During the course of time, various methods have been used to estimate run dynamics and passage counts at point census stations. The simplest methods are based on “connect-the-dots” type of linear interpolation (Gewin and WanHatten 2005, Johnson et al. 2007) but such approach requires an observation before and after the missing time point and thus missing tails cannot be estimated. If it is assumed that a proportion passing the site on a given date is constant over the years then estimation of missing tails is possible using the methods proposed by e.g. Van Alen (2000) but considering the variability, for example, in annual environmental conditions (e.g. Erkinaro et al. 1998; Orell et al. 2007; Otero et al. 2014), such assumption is problematic.

Hilborn et al. (1999) used a maximum likelihood method to estimate number of salmon with ground-based stream survey, accounting also for estimate of uncertainty. Su et al. (2001) continued their work with hierarchical Bayesian approach enabling learning from years with more data to those with missing data. Furthermore, Sethi and Bradley (2016) introduced a Bayesian approach to estimate missing passage at weirs with run curve model to account for arrival dynamics and process variation model to describe the observed data. While these studies attempt to account for uncertainty, they do not discuss their model assumptions against biological knowledge. This together with consideration of difference between the observation and model prediction as “noise” instead of proportion unobserved (as e.g. in Sethi and Bradley 2016), may cast some doubt on the biological meaningfulness of the results.

In our study, a Bayesian model framework was introduced for estimating the annual number of Atlantic salmon (*Salmo salar*) smolts passing the video monitoring site at a subarctic River Utsjoki, a tributary of the large River Teno in northern Finland (cf. Erkinaro et al. 2019). The framework consists of sub-processes of departing, travelling and observing. When combined, the processes of departing and travelling form the arrival distribution that describes the timing of the smolt run. Environmental covariates were used as a source of information in the sub-processes and expert knowledge is utilized in providing biologically meaningful model structures and prior distributions. Hierarchical structure was assumed over the study years making it possible to learn from the processes, borrowing strength from data rich years of study to those with missing data. Predictive studies illustrate the model performance in cases with missing data either at the beginning or in the middle of the run. We show that a biologically meaningful passage model can be built without assuming any specific shape for the arrival distribution. Furthermore, we claim that existing expert knowledge about the observability of smolts is a key source of information in estimating the total uncertainty about the annual total passage especially when environmental conditions have a great influence on the probability to observe individuals passing the survey site. Eventually, the resulting estimates describe the logical inference about the smolt passage that is formed by carefully analyzing the structure of the problem and interpreting observed data in the light of expert’s existing knowledge about migratory behavior and accuracy of video counting.

## Materials and Methods

### Data

River Utsjoki (drainage area 1652 km^2^) is a tributary of the large River Teno system at the northernmost border between Finland and Norway. Salmon production areas extend c. 60 km upstream in the Utsjoki mainstem as well as c. 30 km along two major tributaries (see Borgstrøm et al. 2010). Each spring, the monitoring system of 8 video cameras is set on the river bed at the main channel under a bridge close to the tributary outlet (Figure 1). The video footage of these cameras provides data on both smolts descending and adults ascending the river during the course of summer. Most of the individuals migrate via the main channel, but occasional video monitoring in the side channels has indicated that a few percentages of the smolts would pass through the sides. The installation and use of cameras is described in detail by Davidsen et al. (2005) and Orell et al. (2007). The camera technology and operation have been similar throughout the monitoring years (2002 - present), but some updates have been done to the data recording system, without affecting comparability of the data. In this study we analyze the smolt count data at the main channel from years 2005-2009 and 2014. These years were chosen to demonstrate the highly variable environmental conditions, especially the strongly varying discharge (Figure 2). The smolt data are aggregated in daily counts of individuals over 61 days of June-July.

**Figure 1:**
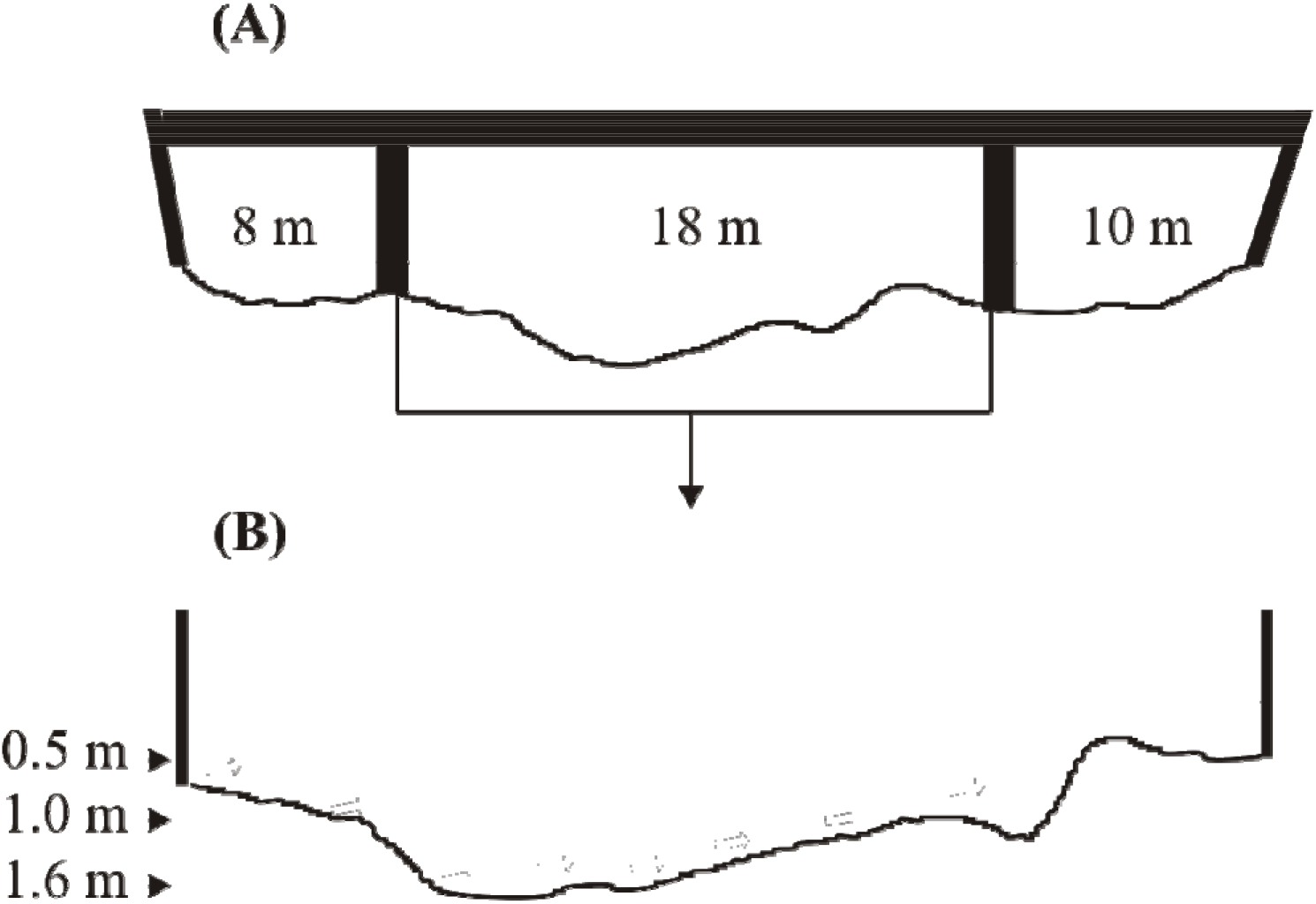
Setup of the video monitoring system at the river bed in Utsjoki. Graph (A) illustrates the profile of the bridge and river bed and graph (B) the river bed in the main channel. Arrows in graph (B) illustrate the position and direction of each of the 8 cameras. Field of view (FoV) of the cameras are: horizontal 78 degrees; vertical 58 degrees.

The datasets on environmental covariates, air temperature and discharge, are aggregated into daily averages. Both covariates are measured near the video site, but conditions may differ between the video site and upper parts of the river where some of the smolts begin their migration from. These datasets are, however, considered as reasonable proxies for the environmental conditions affecting the migration behavior of smolts.

Data from the first 23 days of June in 2005 are missing because the high level of water delayed the installation of the monitoring system. Thus, the missing counts are predicted for those dates. Predictive studies were performed by treating data in 2007 and 2014 as partly missing: For 2007 the first 17% of the run strength and for 2014 the peak +-2 days count was removed from the analysis. These years were chosen to the predictive study as the environmental conditions in those years fell in between the extreme conditions of the overall study years. To evaluate the predictive performance of the model, the model predicted counts for the dates treated as missing were compared with the observed data.

### Expert elicitation as a source of informative priors

Model structure and informative priors were elicited from an expert, who is most familiar with the behavior of salmon smolts and the video monitoring system at Utsjoki. In addition, the expert has comprehensive personal experience on studying smolt migration in other tributaries of the River Teno catchment, as well on other salmon rivers in northern Fennoscandia. The expert also has extensive knowledge about various visual monitoring methods like video monitoring and snorkelling, and has studied their possibilities and restrictions.

The expert was asked to base his views on his overall biological knowledge about the behavior of the smolts: when do the smolts potentially make their decision to depart given the temperature, how long their travelling may take from the nearest and furthest locations of origin, and how the environmental conditions affect the proportion of smolts that may pass the video site unobserved. Extra care was taken to ensure that the expert would express his views based on knowledge independent of the data that would be analysed in the model since the avoidance of double usage of the same data is vital for reliable estimation of total uncertainty.

Elicitation was carried out iteratively, first by asking preliminary questions on e.g. the biological process in question, and then providing graphical illustration of the parameters as a function of environmental conditions (as in Figures 7–9). Prior distributions were adjusted until the expert agreed that the illustration is in line with his views. The questions were not defined - or the answers translated – directly to inform about prior distributions of specific parameters as the marginal distributions often lack good real-life interpretation. Instead, the focus was on the visual form of the joint distributions of each sub-process as a function of an environmental covariate, as will later be shown in figures 7–9. The marginal distributions of parameters related to each sub-process were adjusted until the joint distribution corresponded roughly the view of the expert. The direct questions mainly addressed the mean values of the sub-processes and the levels of environmental covariate at which significant changes in the process were expected to take place. Questions elicited from the expert are listed in table 1.

**Table 1.** Questions elicited from the expert

| Question | Expert view |
| --- | --- |
| *Process of departing* |  |
| - At what range of temperature (air temperature at Utsjoki logger) first smolts are considered to begin their migration? | 6-8 °C |
| - What level of temperature would be considered so high that the process of migration not be influenced by further increase in temperature (i.e. logit curve levels out)? | 12 °C |
| *Process of travelling* |  |
| - What is the the maximum travel time (in days) in which a smolt can be considered to arrive at the video site? | 14 days |
| - What is the minimum travel time in which a smolt can be considered to arrive at the video site? | 1-2 days |
| *Process of observing* |  |
| - What is the upper (/lower) limit for the expected probability at which an average smolt is considered to be observed in best (/worst) possible environmental conditions, i.e. at minimum (/maximum) discharge? | 0.9 (0.3) |
| - What is the level of discharge below which the visibility is considered optimal? | 20 m<sup>3</sup>/s |
| - At optimal visibility, what is the range of probability at which the fish are considered to be observed with around 90% of certainty? | 0.75-0.9 |
| - What is the level of discharge at which the visibility is considered to be poor? | 50-60 m<sup>3</sup>/s |
| - At poor visibility, what I the range of probability at which the fish are considered to be observed with around 90% of certainty? | 0.3-0.6 |

### Probability model of the problem

The model is constructed by thinking of four steps, described in detail in the following sections:

1. Process of departing, in which the smolts make the decision to depart while daily temperature affects the probability to depart in a given day;
2. Process of travelling, in which the smolt’s travel time (in days) from the place of departure to the video site is modeled with daily discharge as a covariate;
3. Arrival distribution, which links together the processes of departing and travelling into a timing of the smolt run;
4. Observation process, in which the discharge influences visibility and the probability that an individual smolt passing the site is observed from the video footage.

### Process of departing

An individual smolt’s probability to begin the migration in day *i* of year *y* given that the smolt has not departed yet is considered to depend on the temperature on that day. We assume a logit-normally linear relationship for the smolt’s probability to depart (*p_i,y_*) and temperature *t_i,y_*:

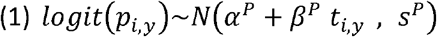

Informative prior distributions are given to parameters *α^P^, β^P^* and *s^P^* according to expert view (see tables 1 and 2).

**Table 2.**
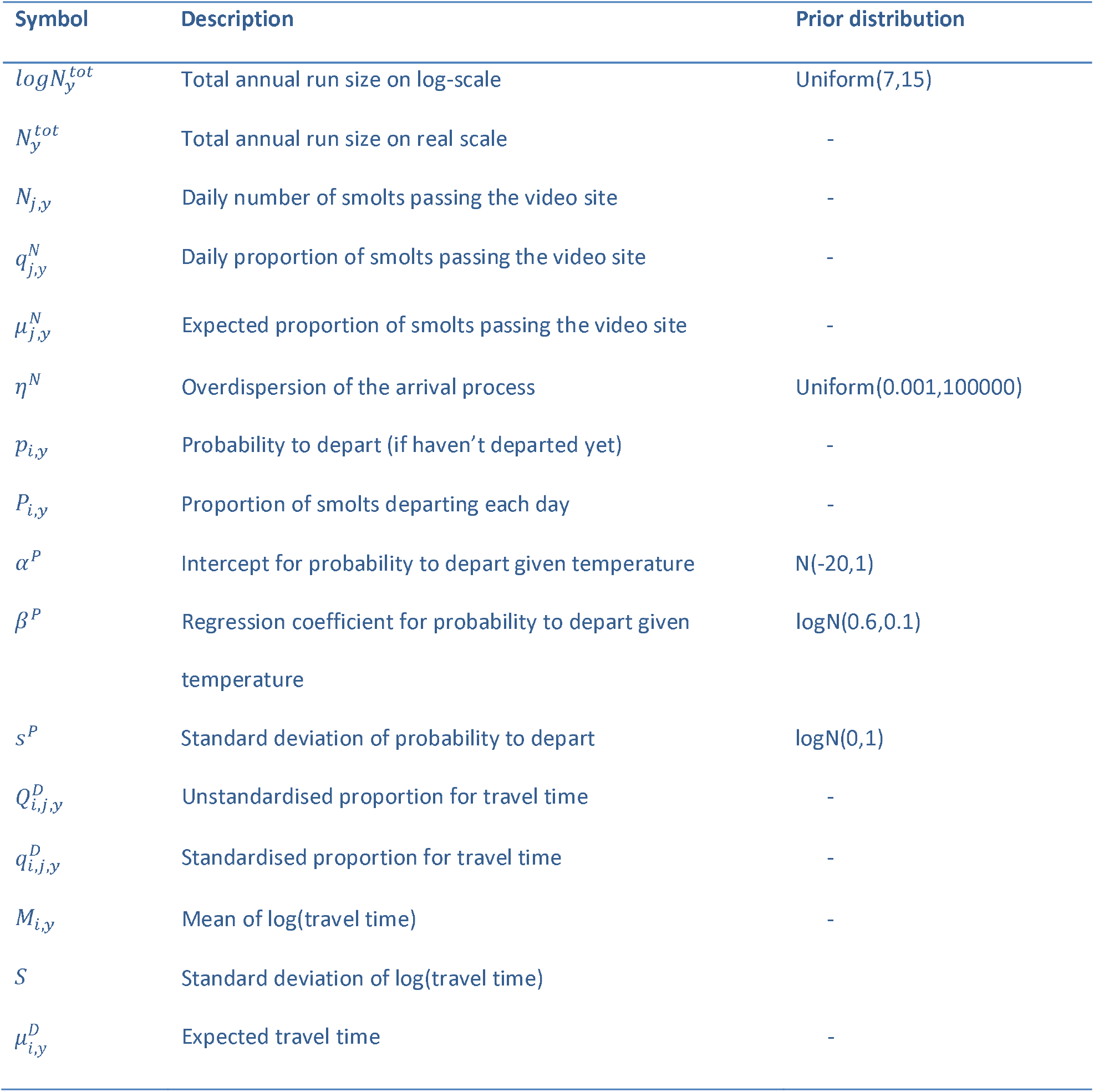
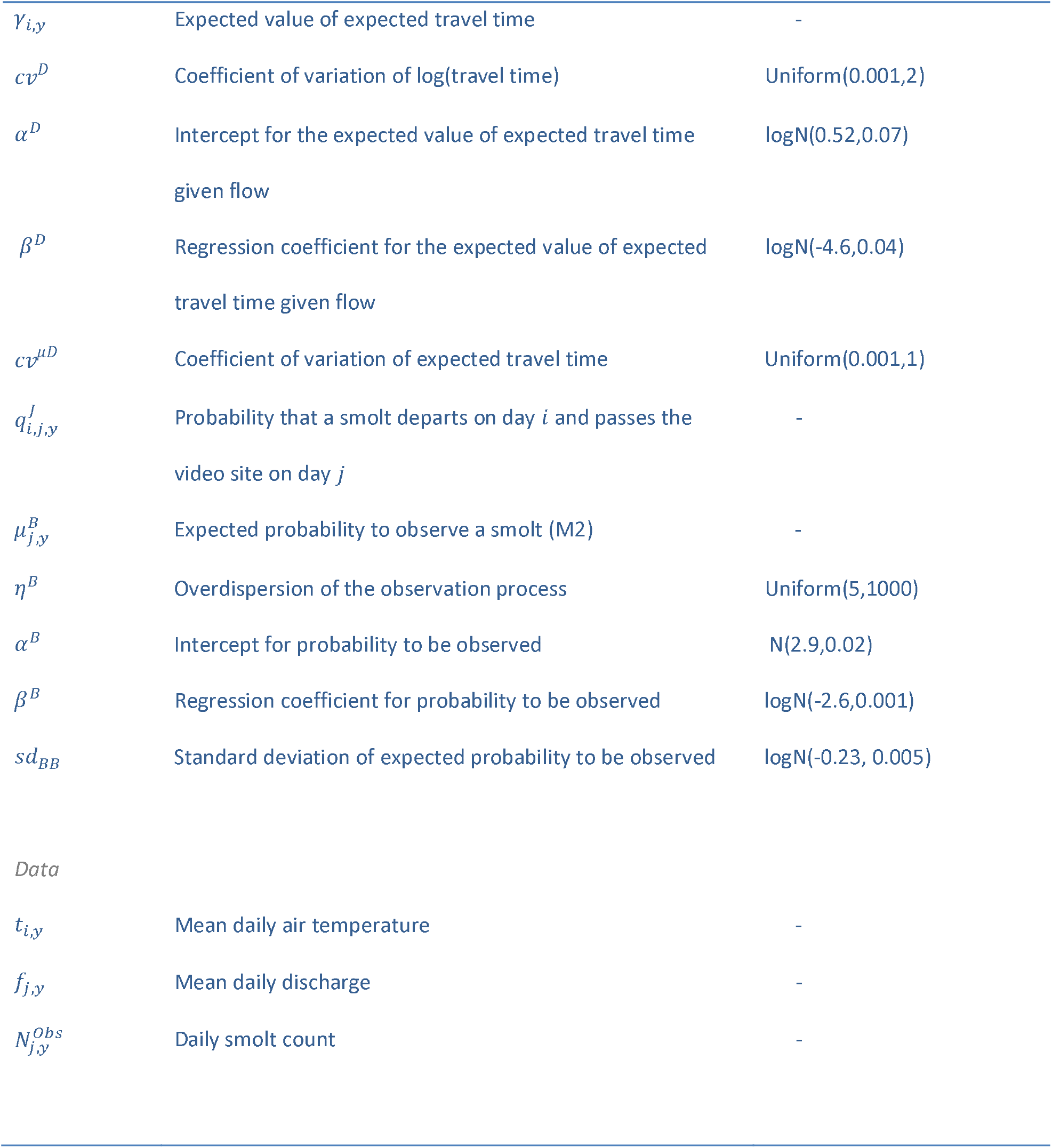
List of symbols, descriptions and prior distributions

The daily probabilities of departing (given those individuals have not departed yet; *p_i,y_*) can be transformed into expected proportions of the total smolt run that depart each day (*P_i,y_*) as

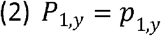

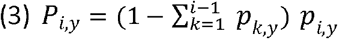

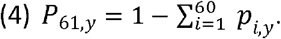

### Process of travelling

According to the expert view, each smolt should arrive at the video site within 14 days after departing. Thus we assume that the smolts that depart on day *i* pass the video site in one of the next 14 days according to discretized distribution function of lognormal distribution (Schwarz & Dempson 1994, Mäntyniemi & Romakkaniemi 2002):

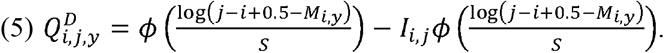

Here *ϕ* is the cumulative density function of the standard normal distribution, *M_i,y_* is the mean of log(travel time) to video site of smolts that depart on day *i*, and *S* is the standard deviation of log(travel time). Thus, 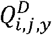 is the conditional probability that a smolt that departed on day *i* will arrive on day *j*, when *j* goes from *i* to *i* + 13.

To ensure that all smolts arrive within 14 days of departing, 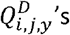 are standardized:

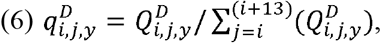

and thus 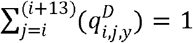.

It is further considered that increasing discharge increases the speed at which the smolts travel, shortening their travel time. The expected travel time on real scale 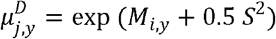 is assumed to follow a lognormal distribution with expected value *γ_i,y_* and coefficient of variation *cv^D^* and it is assumed to depend on the discharge on the day of departure (*f_i,y_*):

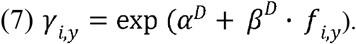

Again, informative prior distributions are given for parameters *α^D^, β^D^* and *cv^μD^* according to expert view (see tables 1 and 2).

### Arrival distribution

By combining the processes of departing and travelling, we achieve joint probabilities that a smolt departs on day *i* and passes the video site on day *j*:

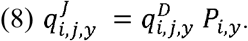

The expected proportions of smolts (out of the total run size) passing the video site each day can then be found by marginalizing over the day of departure:

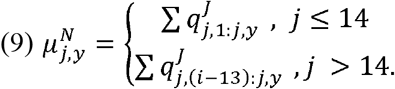

Furthermore, we give a uniform prior distribution on log scale for the total annual size of the smolt run passing the video site:

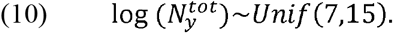

Parameterisation on the log-scale allows for a wide, minimally informative prior distribution for the annual run size. One could also use a more biologically realistic option for estimating the annual abundance based on adult escapement or on expert evaluated smolt production capacity of the river (e.g. Uusitalo et al. 2005). Because the results of this analysis are going to be used in the future research as building block of a sequential modelling framework for the total life cycle of Utsjoki salmon (similarly as in Michielsens et al. 2008), a minimally informative prior distribution for annual production is needed. This ensures that the information flowing from the full life cycle model will be used only once.

The annual run size is assumed to be Dirichlet-multinomially distributed over the 61 days of June-July. For computational simplicity, we follow Mäntyniemi et al. (2015) and approximate multinomial distribution as

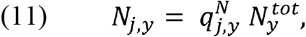

where 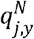 is the proportion of smolts passing the video site in day *j* and 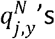 follow a Dirichlet distribution parameterized with the expected proportions of smolts passing the video site each day 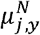 and overdispersion *η^N^*:

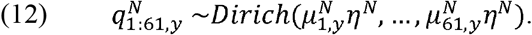

To ease the computation we approximate the Dirichlet-distribution with a set of lognormal distributions (see appendix A for details).

The annual proportional prior distributions for the the smolt run over 61 days of June-July are illustrated in figure 3. These priors are based on the model structure and data on environmental covariates (daily mean air temperature and discharge).

### Observation process

In previous sections we have covered the processes assumed to affect the timing of the smolt passage at the video site. To estimate the total uncertainty in the smolt run size, the process of observing smolts from the video footage needs to be included. It seems natural to assume that some of the individuals pass the site unobserved, and thus this proportion must be accounted for. We introduce an observation process that builds on expert view on how high the probability to observe a smolt may be in excellent vs. poor conditions on visibility and how the discharge is considered to affect this probability. The number of smolts observed follows a Beta-binomial distribution:

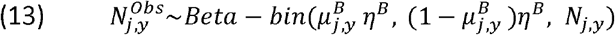

Here *N_j,y_* is the total daily number of smolts passing the video site and the probability to observe a smolt follows a 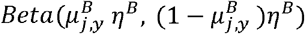 distribution, where 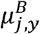 is the expected probability for a smolt to be observed and *η^B^* is the overdispersion parameter. Overdispersion in this context is thought to arise from the schooling behavior of smolts. Groups of smolts can be either detected or missed, which increases the variance of the observation process (Mäntyniemi & Romakkaniemi 2002, Linden & Mäntyniemi 2011, Mäntyniemi et al. 2015). The expected value of the process is linked to discharge following the expert view that in very good visibility (low discharge) at maximum 90% of the smolts are observed. As the discharge increases, the visibility decreases, and the volume of water to be covered by the video monitoring increases. Therefore, according to the expert, observation probability decreases gradually to a minimum of 30%. The expected probability 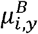 is considered to follow logit-normally linear relationship on the interval [0.3,0.9]:

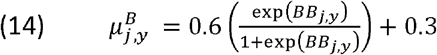

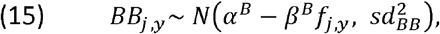

where *α^B^, β^B^* and *sd_BB_* have prior distributions according to the expert view (Tables 1 and 2).

### Bayesian inference and sensitivity analysis

Bayesian analysis that is not sensitive to prior information simply cannot exist. The most influential component of the inference just gradually shifts from prior information about the parameters towards prior information about the data collection process as more data accumulates. From the inferential point of view, both extremes and anything between are equally valid: the resulting posterior shows what our logical state of mind should be after interpreting the data. Sometimes the state of mind is that we learned nothing new about all or some of the parameters.

However, a commonly held belief is that large quantities of data “override” the prior knowledge and thus decrease the sensitivity of the resulting posterior distribution to prior information. Here we show why this is not the case. The key fact is that it is not the data itself that changes the prior, or leaves it unchanged. Prior information becomes updated by interpreting data, and that interpretation is based on prior information. This can be seen from the Bayes’ rule

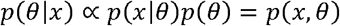

which states that the posterior distribution of parameters (*θ*) is proportional to the joint probability distribution of parameters and potentially observable data (*x*). This distribution summarizes what is known about the entire problem before obtaining the observed data values.

All the joint probabilities *p*(*x, θ*) have a Bayesian interpretation of being measures of degree of belief. In principle, the person whose belief is being measured could state these joint probabilities directly. However, in practice most people find it easier to construct their joint beliefs by factorizing their thinking between what they knew about the parameters (*p*(*θ*)) and how they would use their existing knowledge to predict data if they knew the parameter values (*p*(*x|θ*) (More elaborate factorization can also be used, as demonstrated in previous sections).

The conditional probabilities *p*(*x|θ*) describe the degrees of belief about the link between parameters and data, providing a predetermined interpretation of any potential data values. Once the values are obtained, the posterior distribution (up to a constant) is found by keeping only those joint probabilities that had been pre-assigned to values that became observed. In other words, the observed data objectively selects the subjective joint prior degrees of belief that will be kept to form the posterior distribution.

### Why sensitivity analysis?

As shown above, the results of any Bayesian model are always guaranteed to be sensitive to some aspects of prior knowledge. The role of a sensitivity analysis is to provide information about the relative importance of these different pieces of prior information. Some idea of the relative roles of the information sources can be obtained simply by comparing the priors and posteriors: if the posterior distribution of a parameter is (almost) identical to its prior, then we can conclude that compared to our existing knowledge, our interpretation of the data did not provide new information. This is a case where a sensitivity analysis can reveal deeper insight on the structure of the problem: would our interpretation of data provide at least some new information if we hypothetically had less information about the parameters to begin with? Such a question can be answered by repeating the analysis with priors that express more uncertainty than we actually have.

### Sensitivity study on observability priors

Our preliminary results of the arrival model indicated that parameters *α^B^* and *β^B^* which describe the relationship between the observability of smolts and the discharge, did not update from their priors. In order to gain more in-depth understanding of this result, we performed a sensitivity analysis by pretending that our prior knowledge would be weaker than it actually is: parameters *α^B^* and *β^B^* were given more uncertain prior distributions compared to priors based on expert elicitation (Table 2). More details and results of this study can be found from Appendix B.

### Details on model run

Model covering all the sub-processes was fitted using MCMC sampling with JAGS 4.2.0 software (Just Another Gibbs Sampler; Plummer 2003) and R runjags package (Denwood 2016). Model was run with 2 chains for 7 500 000 iterations which took around 20 days. Burn-in period of 500 000 iterations was removed from the beginning of the run. Convergence was diagnosed using Gelman-Rubin statistics (Gelman and Rubin 1992), ensuring that the upper limit for the potential scale reduction factor was below 1.1 for all parameters of interest. The complete JAGS code for the model structure can be found from Appendix C.

## Results

When compared with the observed smolt counts, annual mean posterior estimates of daily run size indicate that 20 - 43 % of smolts passed the video site unobserved during the years of study (Figure 4). The highest proportion of unobserved was estimated for 2008, median being 43 % and 90 % probability interval (PI) [38 %, 47 %]. This result reflects the high level of discharge in early/midseason in 2008 (Figure 2).

**Figure 2.**
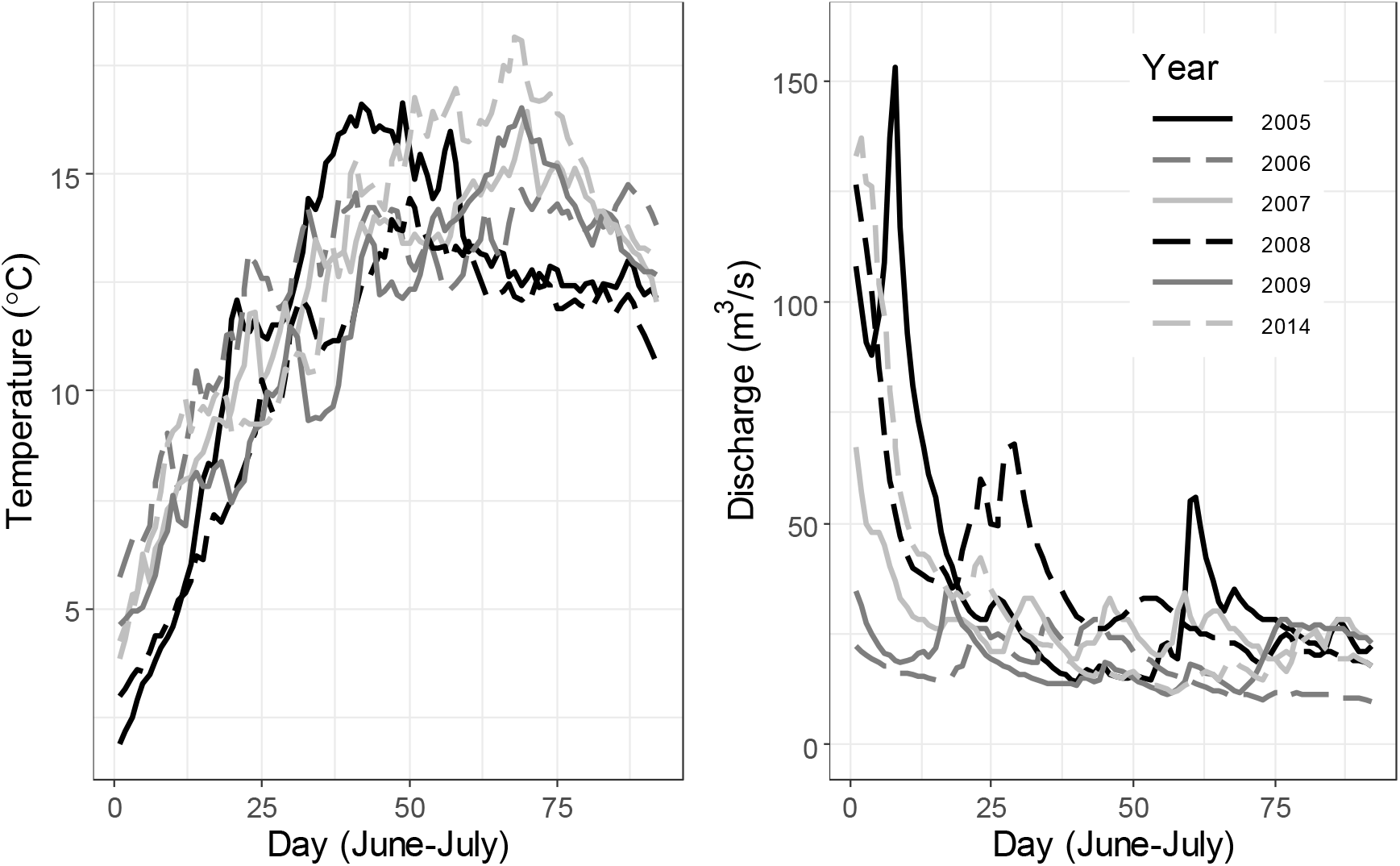
Data on daily mean air temperature and mean discharge at Utsjoki on study years 20052009 and 2014.

**Figure 3.**
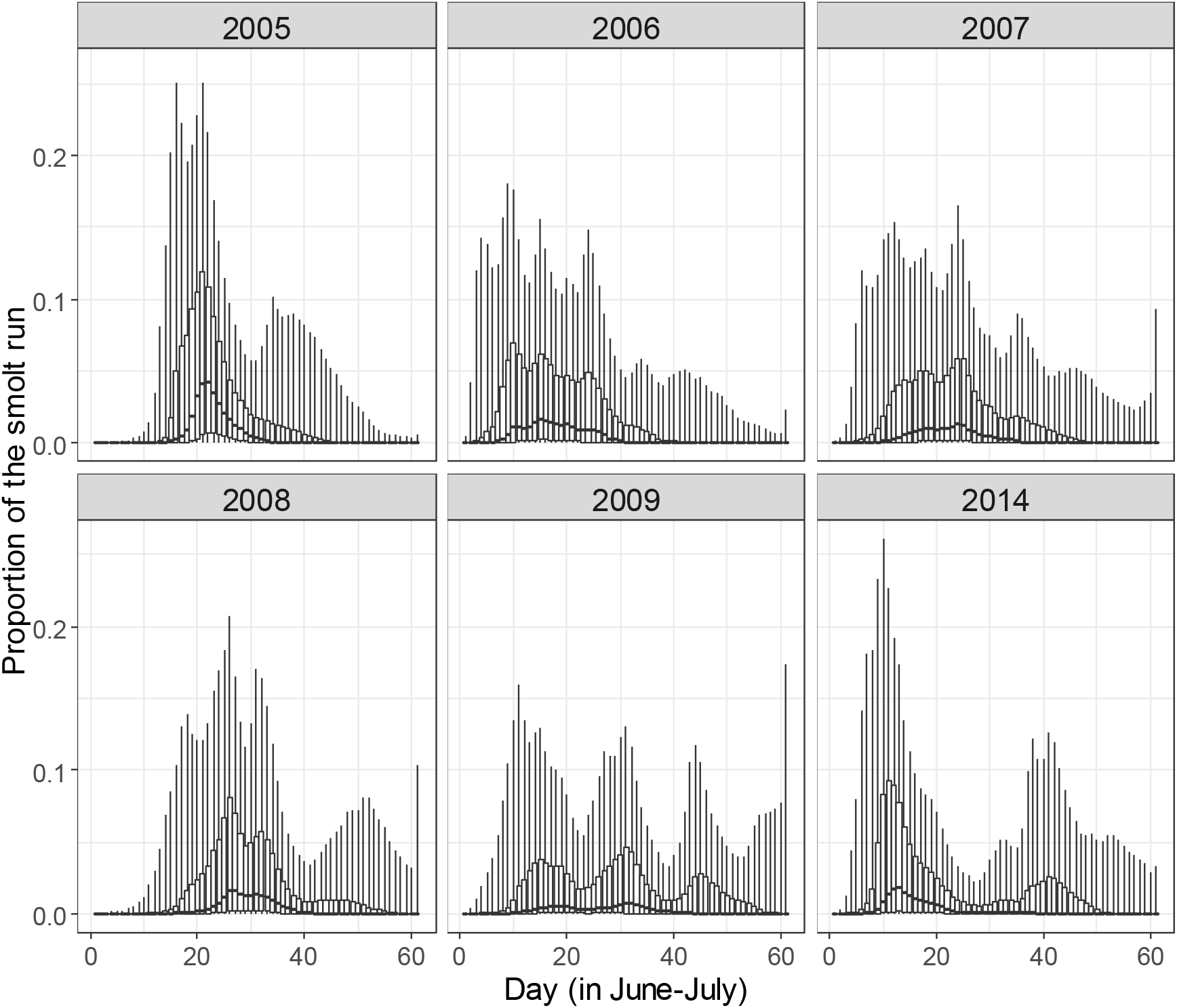
Annual prior distributions for the timing of the smolt run given the environmental covariates (daily mean air temperature and discharge).

**Figure 4.**
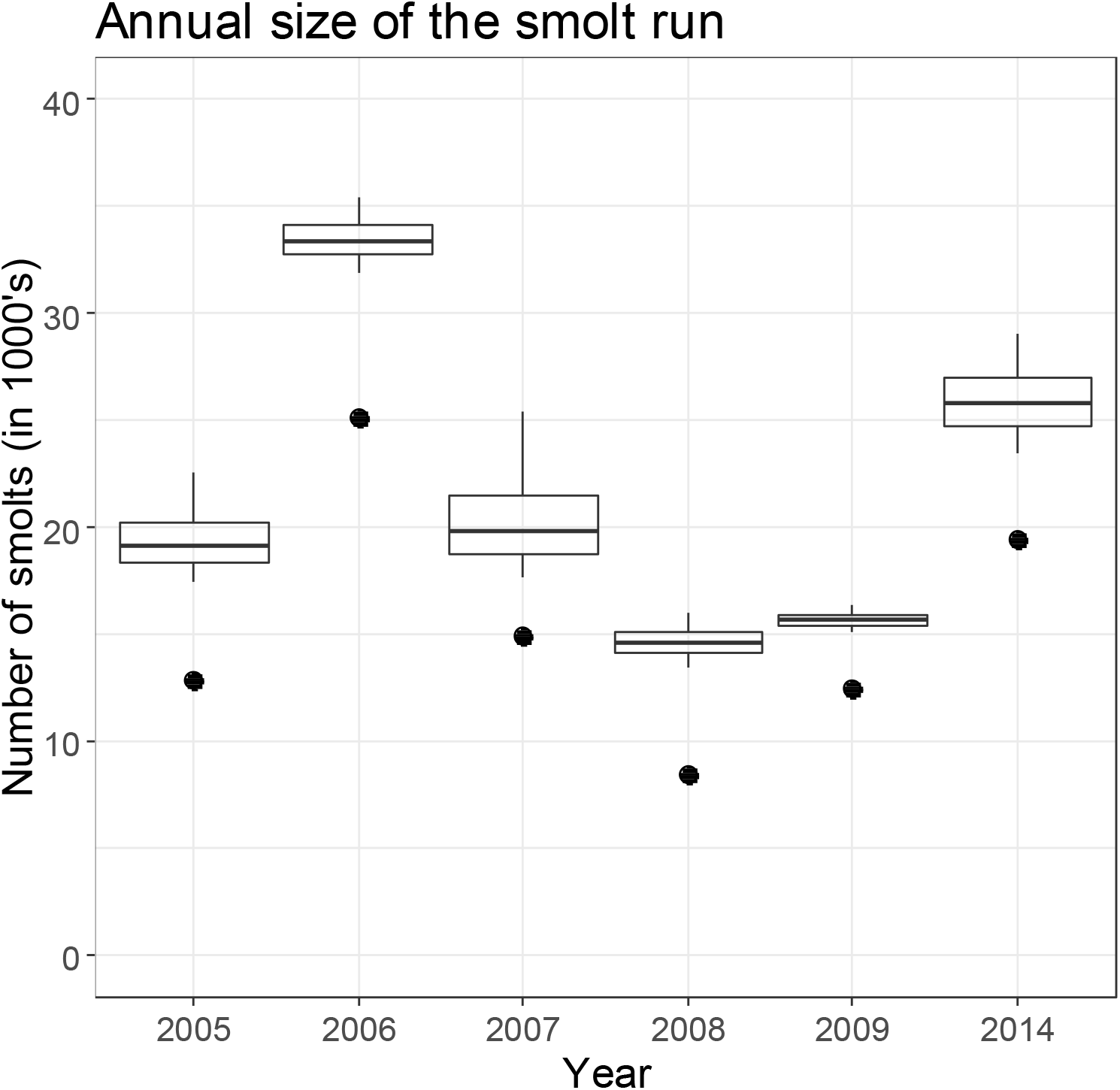
Annual number of smolts passing the video site. Posterior distributions are illustrated with boxplots: middle line is median of the distribution, box covers 50% and whiskers 90% of the probability mass. Count data are illustrated with dots.

Scenarios to study predictive performance of the model (‘first 17% of missing’ -scenario for 2007 and ‘peak day +−2 days missing’ –scenario for 2014) show that daily counts fall in the predicted 90% PI indicating good predictive performance (Figure 5, Table 3). For 2005, when the high level of water delayed the setup of the monitoring system until June 23rd, posterior mean indicates that 10% of the run was missed during this period (90 % PI being [3 %, 23 %] of the run). Because the temperature was below the average until mid-June in that year, it is not likely that the migration would have started very early in the season (Figures 2 and 5).

**Figure 5.**
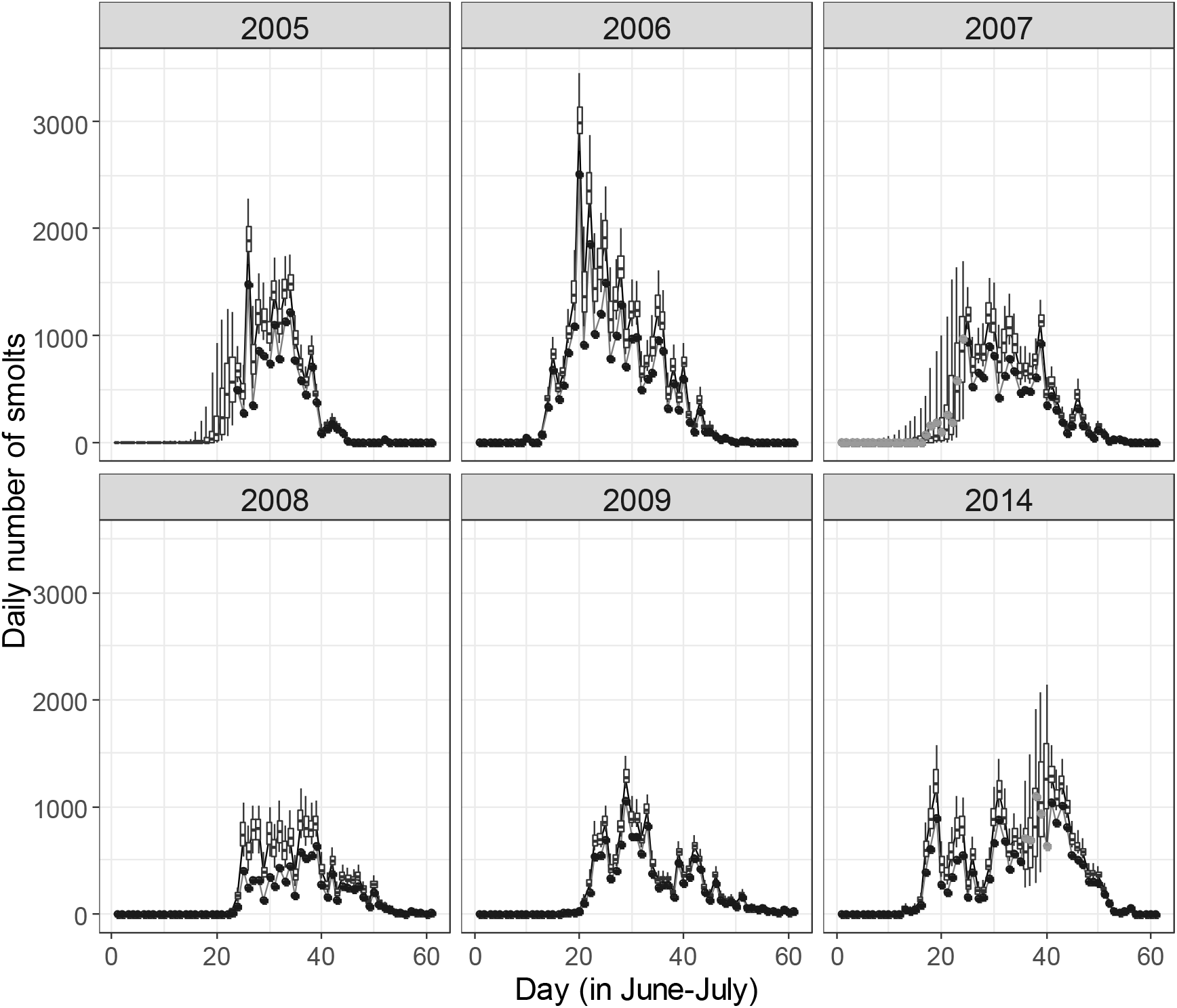
Daily number of smolts that pass the video site in 2005-2009 and 2014. Posterior distributions are illustrated with boxplots: middle line is median of the distribution, box covers 50% and whiskers 90% of the probability mass. Count data are illustrated with dots. Data points included in the study are marked with black and the ones excluded (counts treated as missing) marked with grey.

**Table 3.** Results from the predictive study in 2005 (real missing data), 2007 (‘first 17% missed’ - scenario) and 2014 (‘peak of the migration +−2 days missing’ -scenario). For posterior distributions, mean values and 90% probability intervals are illustrated.

| Year | Sum of missing counts | % of missing counts out of total observed | Number of predicted missing | % of predicted missing out of total |
| --- | --- | --- | --- | --- |
| 2005 | NA | NA | 2122 (457,5159) | 10% (3%,23%) |
| 2007 | 2539 | 17% | 3346 (723,8546) | 15% (4%,33%) |
| 2014 | 4085 | 21% | 4677 (2580,7367) | 18% (11%,26%) |

When comparing marginal posterior distributions with corresponding priors, 7 key parameters out of 12 updated, indicating that the learning does not take place evenly about all parameters (Table 4). The joint posterior distributions of the process of departing as a function of temperature (consisting of parameters *α^P^, β^P^* and *sd^P^*) updates quite heavily from the prior (Figure 6). Updating takes place towards higher temperatures indicating, for example, that 50 % probability to depart is met with 50% certainty around 14 °C compared to a priori belief of this level being reached already at 11 °C.

**Figure 6.**
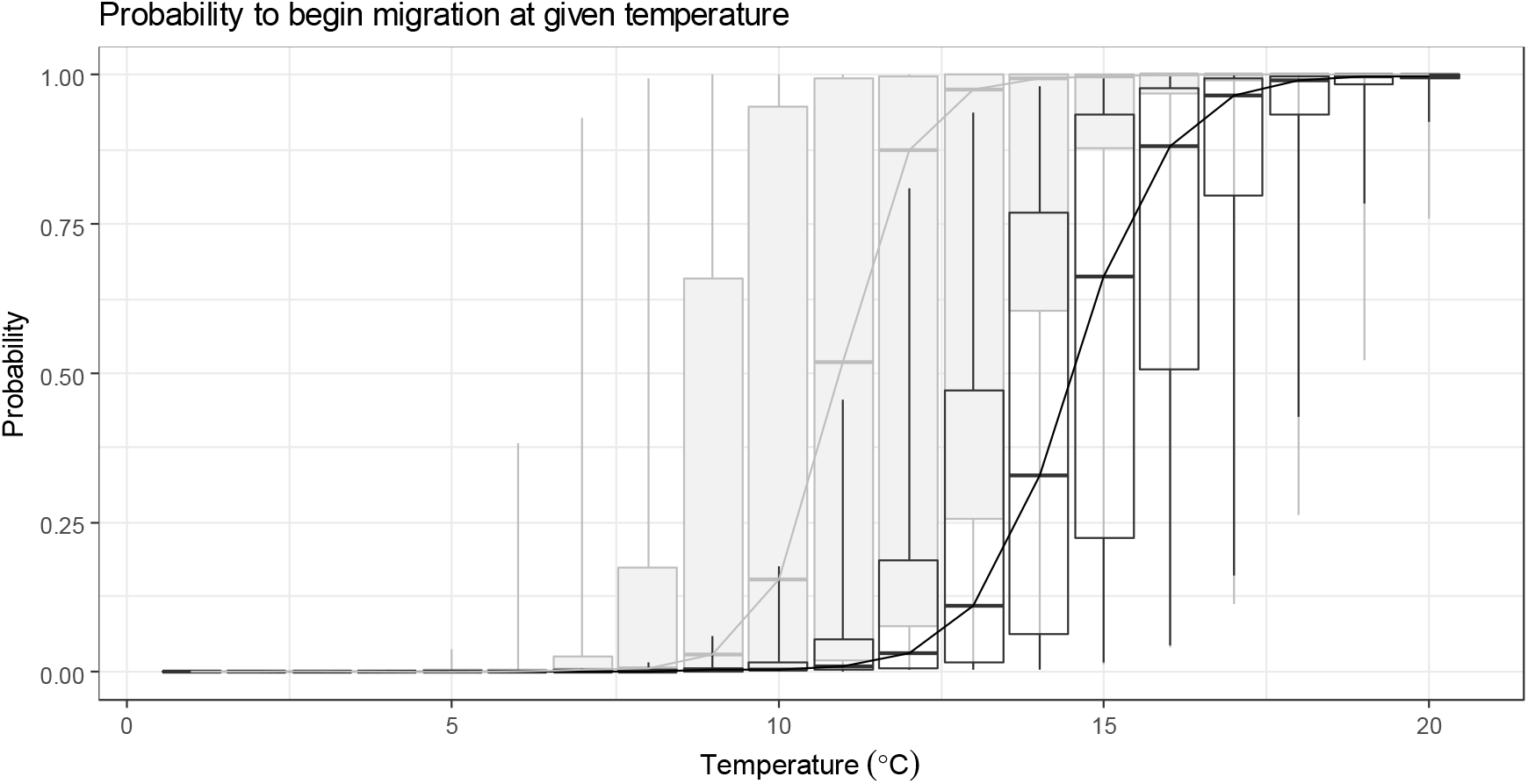
Probability that a smolt begins the migration at given temperature. Priors and posteriors are illustrated with grey and black boxplots, respectively. Middle line in the box illustrates median of the distribution, whereas box covers 50% and whiskers 90% of the probability mass.

**Table 4.** Prior and posterior means and 5 % and 95 % quantiles of key parameters

| Variable | Prior |  |  | Posterior |  |  |
| --- | --- | --- | --- | --- | --- | --- |
|  | mean | 5% | 95% | mean | 5% | 95% |
| $\alpha^P$ | -20.0 | -21.6 | -18.3 | -19.9 | -21.3 | -18.5 |
| $\beta^P$ | 1.92 | 1.09 | 3.09 | 1.37 | 1.25 | 1.49 |
| $s^P$ | 1.65 | 0.20 | 5.15 | 2.89 | 2.36 | 3.54 |
| $\alpha^D$ | 1.74 | 1.09 | 2.61 | 2.63 | 2.21 | 3.09 |
| $\beta^D$ | 0.010 | 0.007 | 0.014 | 0.010 | 0.007 | 0.013 |
| $cv^D$ | 1.00 | 0.11 | 1.90 | 1.54 | 1.01 | 1.95 |
| $cv^{\mu D}$ | 0.50 | 0.05 | 0.95 | 0.22 | 0.02 | 0.55 |
| $\alpha^B$ | 2.90 | 2.69 | 3.11 | 2.91 | 2.70 | 3.11 |
| $\beta^B$ | 0.074 | 0.070 | 0.078 | 0.074 | 0.070 | 0.078 |
| $sd_{BB}$ | 0.80 | 0.71 | 0.89 | 0.81 | 0.72 | 0.90 |
| $\eta^B$ | 505 | 56 | 951 | 408 | 15 | 933 |
| $\eta^N$ | 50092 | 5008 | 95069 | 289 | 208 | 403 |

The posterior cumulative distribution of the expected travel time to the video site (consisting of parameters *α^D^, β^D^, cv^D^* and *cv^μD^*) supported somewhat longer travel times than expected a priori both for low and high discharge (Figure 7). At low discharge (10m ^3^/s), 50 % of smolts were estimated to arrive at the video site within 5 days (posterior median), whereas a priori the same proportion was considered to be met around 3.5 days. At high discharge (100m ^3^/s), 50 % of smolts would arrive by 3 days based on the posterior and by a bit less than 2 days based on the prior. Note that posterior cumulative distributions at these specific levels of discharge were chosen for illustrative purposes and that in practice the model allows any positive value for the discharge.

**Figure 7.**
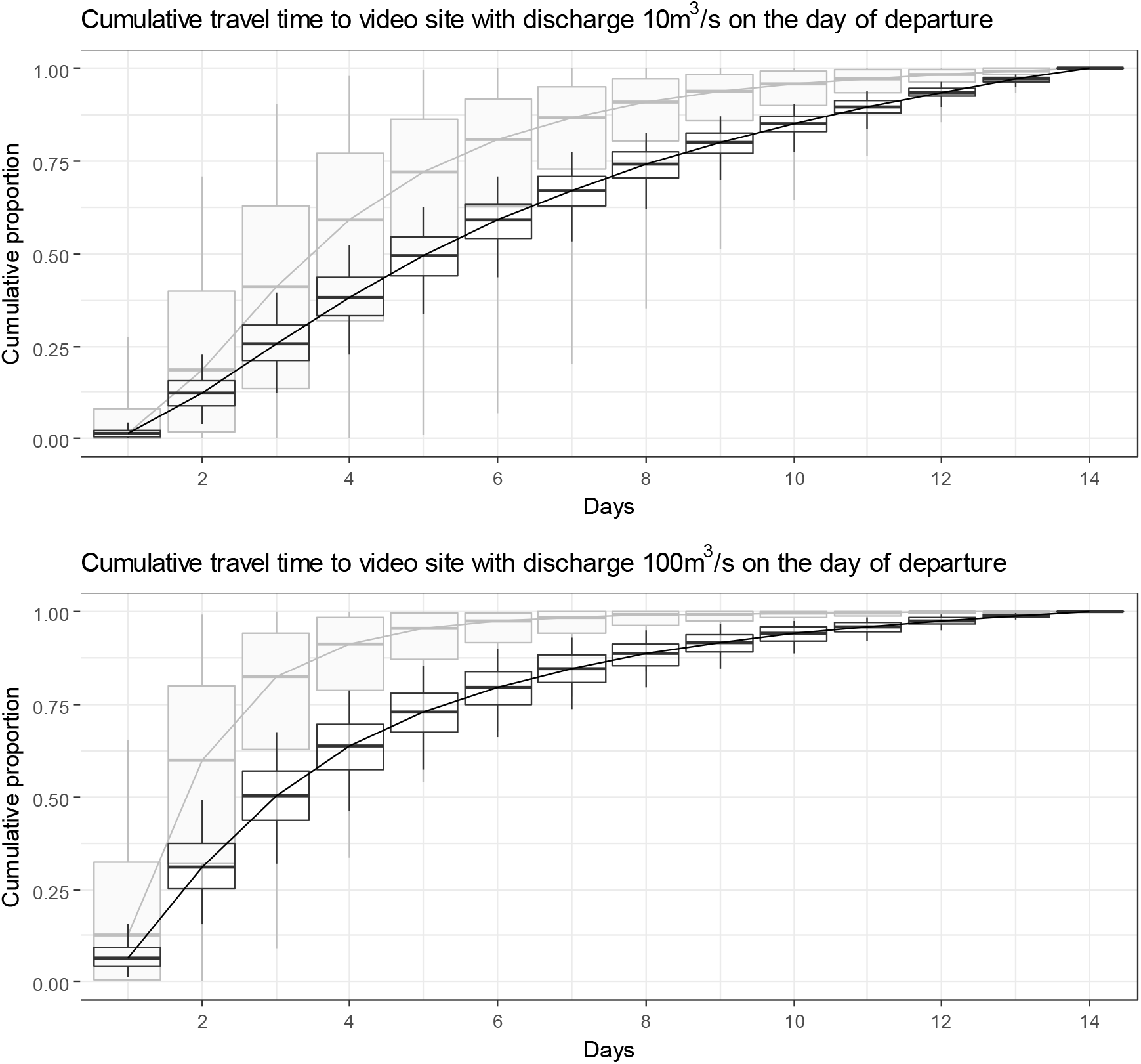
Cumulative distribution of expected travel time in days from the point of departure to the video site. Above: Cumulative distribution at low discharge (10 m^3^/s). Below: Cumulative distribution at high discharge (100 m^3^/s). Priors and posteriors are illustrated with grey and black boxplots, respectively. Middle line in the box illustrates median of the distribution, whereas box covers 50% and whiskers 90% of the probability mass.

Joint priors and posteriors of observability parameters *α^B^, β^B^* and *sd^BB^* are close the same (Figure 8) indicating that in this case the combination of priors and our interpretation of data did not result in further learning about the process of observing. Sensitivity analysis (Appendix B) revealed that if we happened to have more uncertainty about these parameters, we would have clearly learned something new about the role of discharge (*β^B^*) in the observability of smolts. The parameter responsible for the baseline of the observability (*α^B^*) was more sensitive to prior uncertainty about it: no new information after seeing the data could be obtained within the range of parameter values studied.

**Figure 8.**
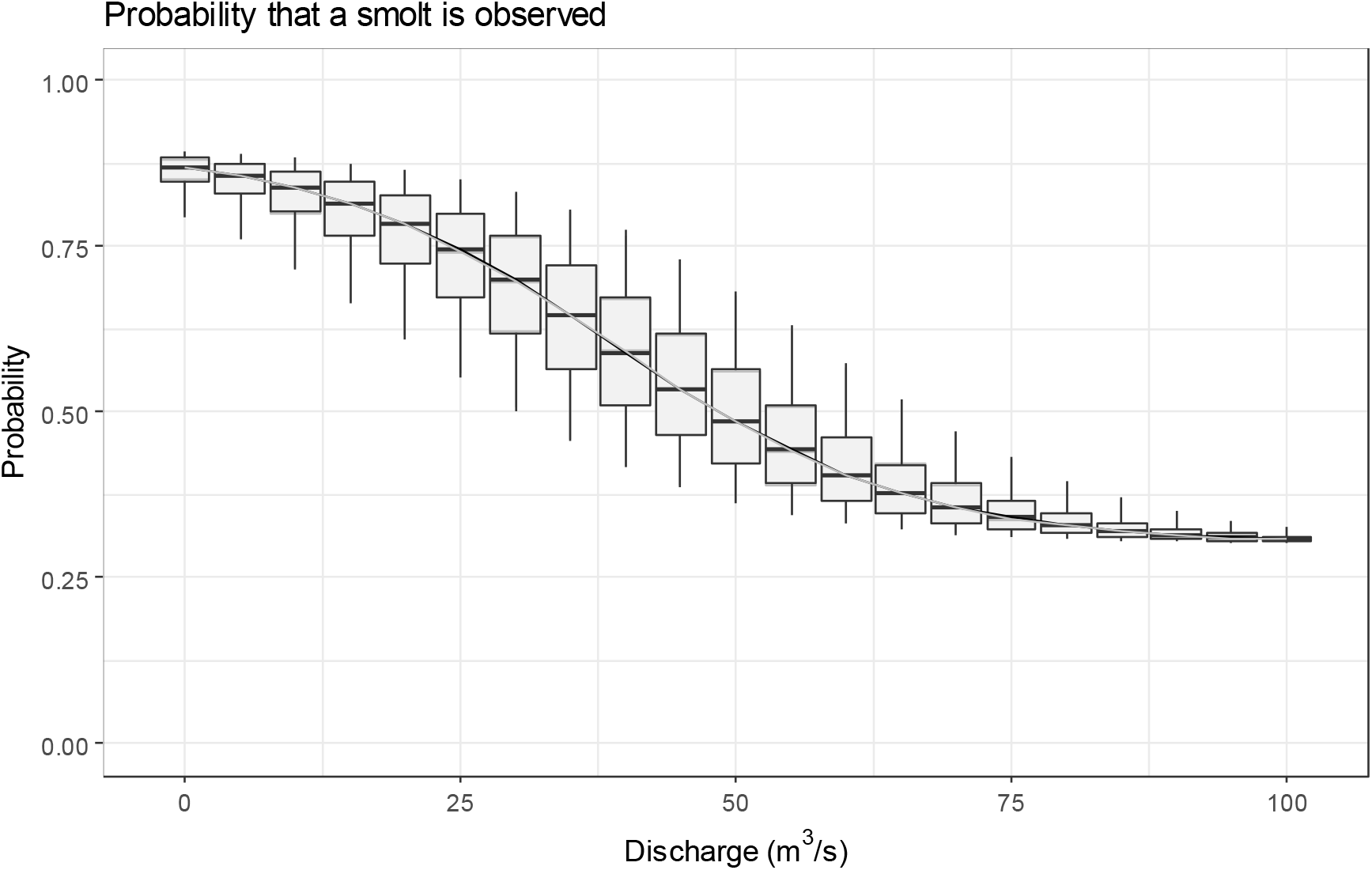
The expected observation probability at given discharge. Priors and posteriors are illustrated with grey and black boxplots, respectively. Middle line in the box illustrates median of the distribution, whereas box covers 50% and whiskers 90% of the probability mass.

## Discussion

In our study, an arrival model for Atlantic salmon smolts was designed by separating the processes of departing, travelling and observing. The processes of departing and travelling together form the arrival distribution, i.e. the timing of the smolt run. This modular structure enabled us to concentrate on biologically meaningful mathematical assumptions related to each sub-process instead of assuming that fish count data as such follows a specific distributional shape. The model framework aimed at high biological realism by accounting for expert knowledge in the model formulation, firstly by including choices for model structures and prior distributions, and secondly by linking environmental covariates to provide information about the timing of the migration. The estimated total uncertainty accounted for the uncertainty in the sub-processes, including the observation process in which variability in discharge is known to have a large impact on the probability to observe passing smolts.

One of the most important advantages of our approach is that the need to assume a specific parametric shape for the arrival distribution is avoided. Past studies (Hilborn et al. 1999, Su et al. 2001, Sethi and Bradley 2016) have assumed unimodal shape (e.g. normal, skew-normal, Student’s t) for the arrival curve. However, when taking a look at the shape of (any) arrival time series it seems clear that such assumption is a tolerable simplification at best (see e.g. figure 5 in this paper, figure 2 in Sethi and Bradley 2016).

Instead of basing the arrival curve on a specific distributional form without biological interpretation, we have based our model structure largely on the expert knowledge. While one can state that the expert may be wrong in his/her views, it can also be argued that replacing the available knowledge with ad-hoc but conventional choices of distribution has even less of a chance to end up with a “true” outcome. The expert elicitation requires ensuring that the experts are able to express their views about the processes independent from the data that will be analysed. This can be problematic at first as often the same experts have the best knowledge about the issues related to conditions at the study area, behavior of the species, details on the data collection and the data itself. We recommend that questions elicited should be designed (similarly as in our study) to target the sub-processes and phenomenon behind the data, not the data itself.

Fitting the elicited information together with the model structure can also be challenging. For example, when we combined the expert views on the observation process with a logistic functional form, very low uncertainty was inevitably assumed at high (>60 m^3^/s) discharge. However, such discharges usually take place only at the beginning of the season when the smolt run has not really started (Orell et al. 2007), and thus the impact of the assumption on the estimated total run size is minor. Regardless of methods chosen on which the model is built upon it is important to keep in mind that the results depend always on the combination of the objective data and subjective model choice.

Earlier studies have paid little attention to the role of observation models in estimating the total uncertainty. Hillborn et al. (1999) distinguished process and observation errors, assuming observer efficiency either as known or unchanging inside the study period. Su et al. (2001) discussed the issue of error in the observation process and mainly concluded that such error can cause convergence problems in sampling. Sethi and Bradly (2016) incorporated uncertainty only on the estimates of missing passage counts, treating the observed counts as true total number of passing without uncertainty. As they assumed normal or negative Binomial models for the observed passage counts, they in fact allowed model predicted passage counts to be either greater or smaller than the observed counts. This in turn can lead to underestimation of the missing counts. We claim that it is necessary to consider the observed count as a minimum estimate for the true number especially if the back-and-forth movement of individuals is negligible. Along with this notion the need for a realistic observation process becomes obvious. Depending on the approach, the estimate of total annual passage can be very different. As Hilborn et al. (1999) put it, ‘when we admit uncertainty in observer efficiency we become much less certain about the actual escapement’.

The sensitivity analysis showed that in our case video counts could provide information about the effect of discharge on the observability. This can be explained by the fact that the daily video counts are thought to depend on the number of smolts migrating and their observability, both of which are thought to depend on daily varying environmental covariates. The daily number of migrating smolts is controlled by temperature and discharge -related parameters, but the model fit to observed counts can be further improved by changing the observability according to discharge. Interestingly, the sensitivity analysis showed that the interpretation of data favors the same parameter values than the expert’s prior. However, even though the signal from the data agrees with the expert’s existing knowledge, the signal is weak compared to expert’s already existing certainty. Therefore the certainty does not notably increase. The sensitivity analysis also revealed that in our case the general level of the observability of smolts is highly dependent on the existing expert knowledge: very little or no new information could be inferred from the video counts. This underlines the need for rigorous expert elicitation.

If the purpose of the model would have been to replicate the observed data set with as few parameters as possible, then the sensitivity of the results to the observability priors would be a sign of overparameterisation: these parameters could be the first to be dropped from the model. The concept of population size would probably be unnecessary too for such a purpose. However, our goal is not to find the most parsimonious model for data, which would be the purpose of information theory. Instead, we are dealing with an inverse problem: given the data and our existing knowledge about migration and observation processes, what can we realistically say about the population size? Within this approach the sensitivity is not of consequence when it comes to simplifying the model. As shown earlier, any Bayesian model is completely made of prior knowledge and always sensitive to it in one way or another. Any model simplification should be based on biological plausibility, rather than on the ability of the model to replicate data (Kuparinen et al. 2012). Using information theoretic arguments to simplify a model meant for solving inverse problems can lead to underestimation of actual uncertainty (Kuparinen et al. 2012) which causes underestimation of the value of any new information that could be potentially gathered (Mäntyniemi et al. 2009).

Sethi and Bradley (2016) fitted datasets in their study using several combinations of arrival and process error models. They utilized information-theoretic model selection (DIC, Spiegelhalter et al. 2002) to find out the best fitting model for each dataset. However, no discussion took place whether some of these models would be biologically more reasonable than another. We argue that the model selection should primarily be based on biological realism and in case biological model uncertainty is needed to be tackled, Bayesian model averaging (BMA, Hoeting et al. 1999) should be preferred over DIC or other information criteria.

Su et al. (2001) used hierarchical structure in their study, assuming that years in which timing of the run is well known can be used for passing information to years with missing data. We argue that learning should rather take place over the (biological) process, assuming exchangeability over year specific parameters of those processes. Our model structure can also be considered as hierarchical, although we simply assume that the underlying processes are the same from year to year allowing between- and within-years variation. If there were a need for a meta-analysis of similar studies over several stocks of the same species, exchangeability in biological parameters could be considered between the stocks. Such approach could also allow meta-analysis of data from different types of monitoring systems if detailed observation processes were tailored for each system and exchangeability was considered as reasonable assumption for the biological processes.

Our model framework could be fine-tuned and developed further. In an ideal situation, environmental covariates would be available from upstream locations from where the smolts originate. Data on air temperature would be replaced with water temperature to better explain the physiological processes the smolts encounter. The total annual estimates could be improved by including individuals that migrate via side channels. One year of video data from the side channels is available but larger set of data would be needed to evaluate the effect of varying discharge on the proportion of smolts passing via the main vs. side channels. To increase the realism even more, the schooling behavior of smolts could be accounted for in more detail than done here. Salmon smolts have a tendency to school during their migration (e.g. Bakshtanskiy et al. 1988, Davidsen et al. 2005) and this creates more variation in the monitoring data compared to independent movement (Mäntyniemi and Romakkaniemi 2002, Mäntyniemi et al. 2015). Schooling behavior could also be linked with the process of travelling as the passing schools of fish have been suggested to stimulate migratory behavior in the remaining smolts (e.g. Hansen and Jonsson, 1985). Schooling also affects the observation process as schools are easier to spot than individual smolts, and on the other hand, it can be difficult to have an exact count of individuals from a large school. For Utsjoki video survey, daily average school sizes of smolts are recorded and incorporating those into the model framework is an interesting topic for future research.

Despite the above possibilities for further development of the modeling framework, we claim that our model already contains the elements of realism that are required to provide credible estimates of the total annual smolt run. This approach can be applied to similar monitoring surveys where point-checking the numbers of migrating fish (Trépanier et al. 1996), by e.g. sonar, trap, or weir is used for estimating run sizes. As far as we know, environmental covariates have not been included so far to estimate both the dynamics of passage and the observation process. Our model framework also indicated a good predictive performance for missing dates both at the beginning and in the middle of the run. Furthermore, the results from this study will act as a single step of sequential model framework that covers the total life cycle of multiple populations in the River Teno complex (Vähä et al. 2017; Erkinaro et al. 2019), similarly as the system designed for Baltic salmon stocks (Michielsens et al. 2008).

## Acknowledgements

We thank the numerous field workers enabling the yearly monitoring programme and smolt data collection at the River Utsjoki. We also acknowledge the crucial technical knowhow by Jorma Kuusela in running the annual monitoring and the instrumental role of Dr. Martin Svenning and Anders Lamberg in early phases of the River Utsjoki monitoring programme. Funding for this study was provided by the Academy of Finland (project No. 286334).

## Appendix A: Lognormal approximation for Dirichlet-distribution

Let’s consider a set of Dirichlet-distributed parameters:

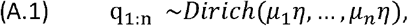

Where *μ_i_* is the expected probability for slot *i* and *η* is the overdispersion parameter. Such Dirichlet-distribution can be approximated with a set of lognormally distributed *z_i_* by assuming

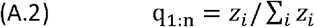

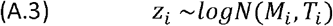

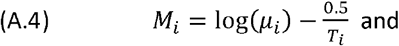

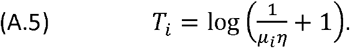

## Appendix B: Sensitivity study on priors of the observation process

The expert elicited priors for the parameters of the observation process (Figure 8) did not update when the model was fitted to video counts. In order to study reasons behind this finding we made another run with the model with more uncertain prior distributions for parameters *α^B^* and *β^B^*. Parameter *α^B^* was set to follow N(2.9,l) and parameter *β^B^* logN(−2.6,1) distribution, respectively. Other prior distributions and model assumptions remained the same as described in the methods section. Model was run 5 100 000 iterations at which point all parameters of interest had converged according to Gelman-Rubin shrink factor.

Unlike in the model run with expert elicited priors, the sensitivity model run with wider prior distributions resulted with update for observability parameters (Figure B1). Parameter *α^B^* updates only slightly, and most of the learning takes place for parameter *β^B^*. In comparison, in the model run with expert elicited priors, posteriors of these parameters were identical to the prior distributions (Figure 8 in Results section). The joint distributions of priors/posteriors of parameters *α^B^, β^B^* and *sd^BB^* as a function of discharge (Figure B2) show that learning takes place mainly at the lower ends of the observation probability.

Joint posteriors from the model run with expert elicited priors and sensitivity model run are compared in figure B3. The difference in the joint posteriors mainly takes place as increased uncertainty on higher probabilities to observe smolts, whereas the lower limits of the distributions are relatively close between the models.

Figure B4 illustrates the impact the prior choice on observability has on the posterior estimates on the annual total number of smolts. The posteriors are close the same at the upper end of the distributions, whereas at the lower ends of the distributions the estimates of the sensitivity model support smaller annual abundances than the estimates from the run with expert elicited priors (Figure B4). We learn that with the current model structure, the video survey provides mainly information about the minimum level of observability (through realized discharges), but that expert knowledge is vital in understanding the maximum limits of observability. Posterior distributions of parameters related to processes of departing and travelling were identical with the estimates of the model run with expert elicited priors on observation process (Figures 6 and 7 in the Results section).

Scatter plots of the posterior samples between the annual total number of smolts and parameters of the observation process, *α^B^* and *β^B^* (Figures B5 and B6), give us more insight on the model behavior. The interpretation of these parameters is that the *α^B^* sets up the base level for the observation probability and *β^B^* defines on how quickly the probability decreases as the discharge increases. The posteriors on total smolt abundance seem to be more sensitive on the choice of prior on *α^B^* than for choice of prior on *β^B^*, this can be seen especially from the samples of the sensitivity model run (Figures B5 and B6). On the other hand, observed data has more power to update parameter *β^B^* than *α^B^* (Figure B1). Correlation between *α^B^* and *β^B^* balances the joint distribution in the sensitivity model run, as with high values of *α^B^* values of – *β^B^* decrease (Figure B7).

**Figure B1.**
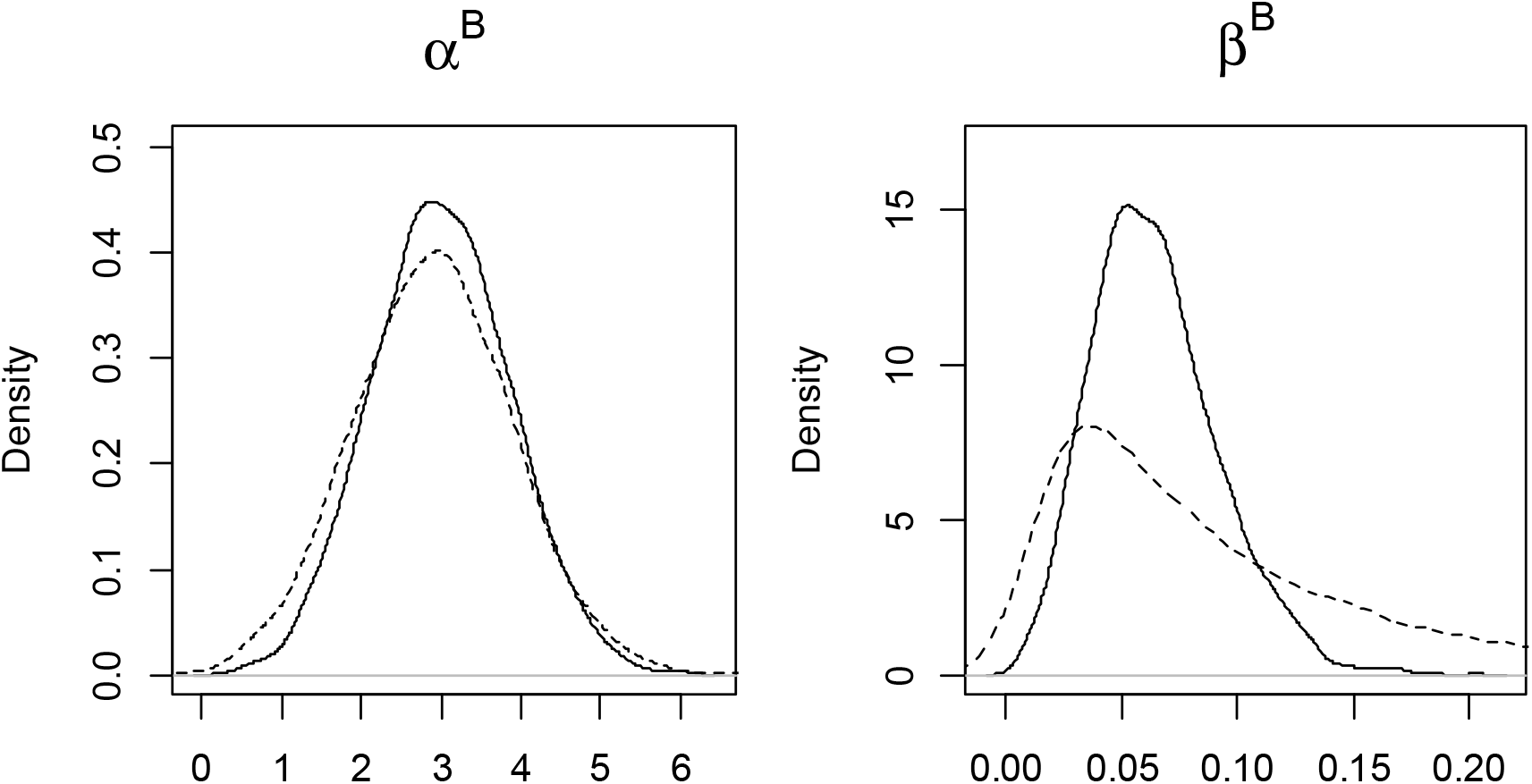
Marginal densities of parameters *α^B^* and *β^B^* in the sensitivity model run. Prior distributions are illustrated with dotted and posterior distributions with solid lines.

**Figure B2.**
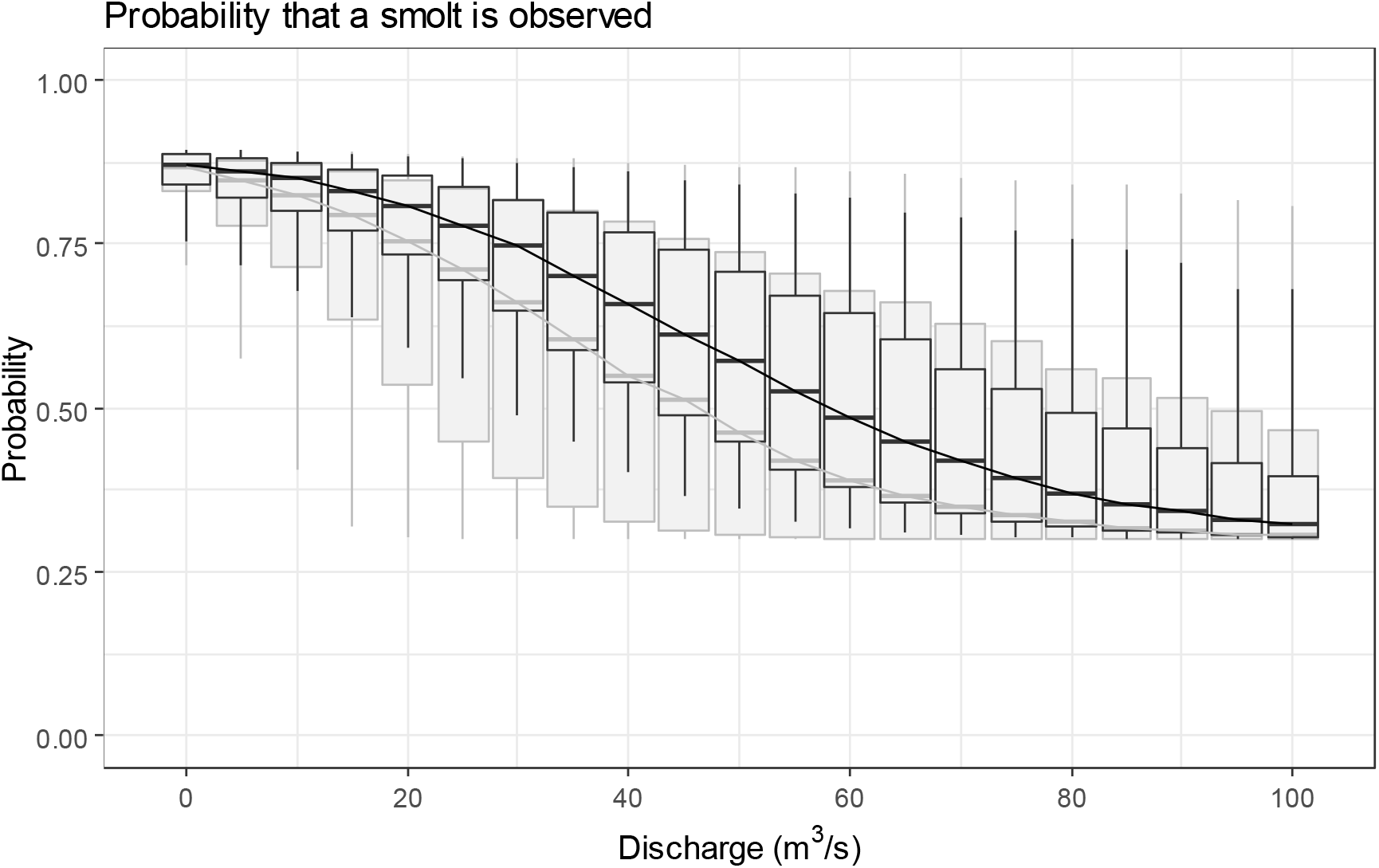
The expected observation probability at given discharge in sensitivity model run. Priors and posteriors are illustrated with grey and black boxplots, respectively.

**Figure B3.**
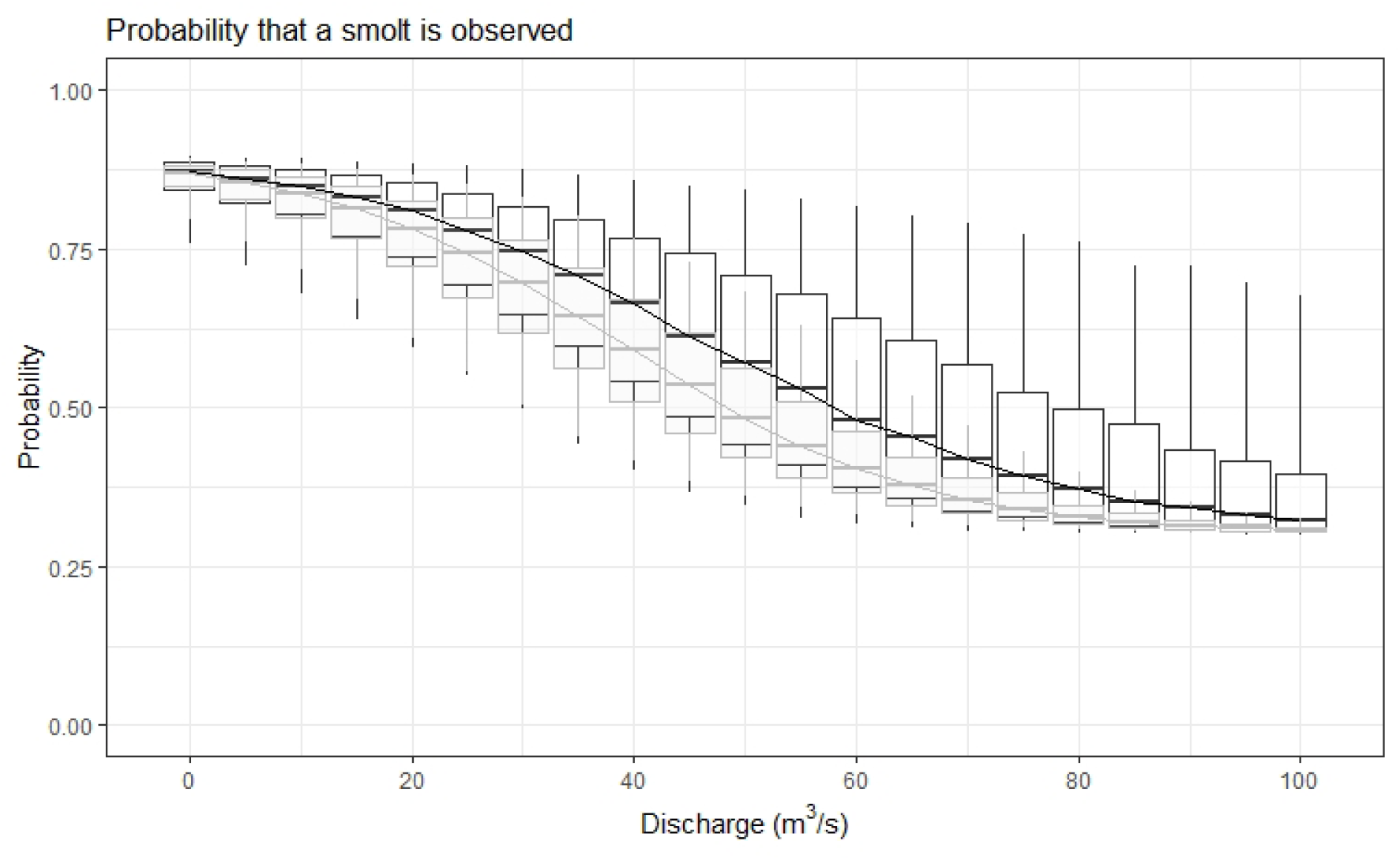
The expected observation posterior probability at given discharge in model run with expert elicited priors (grey box plots) and sensitivity model run (black boxplots).

**Figure B4.**
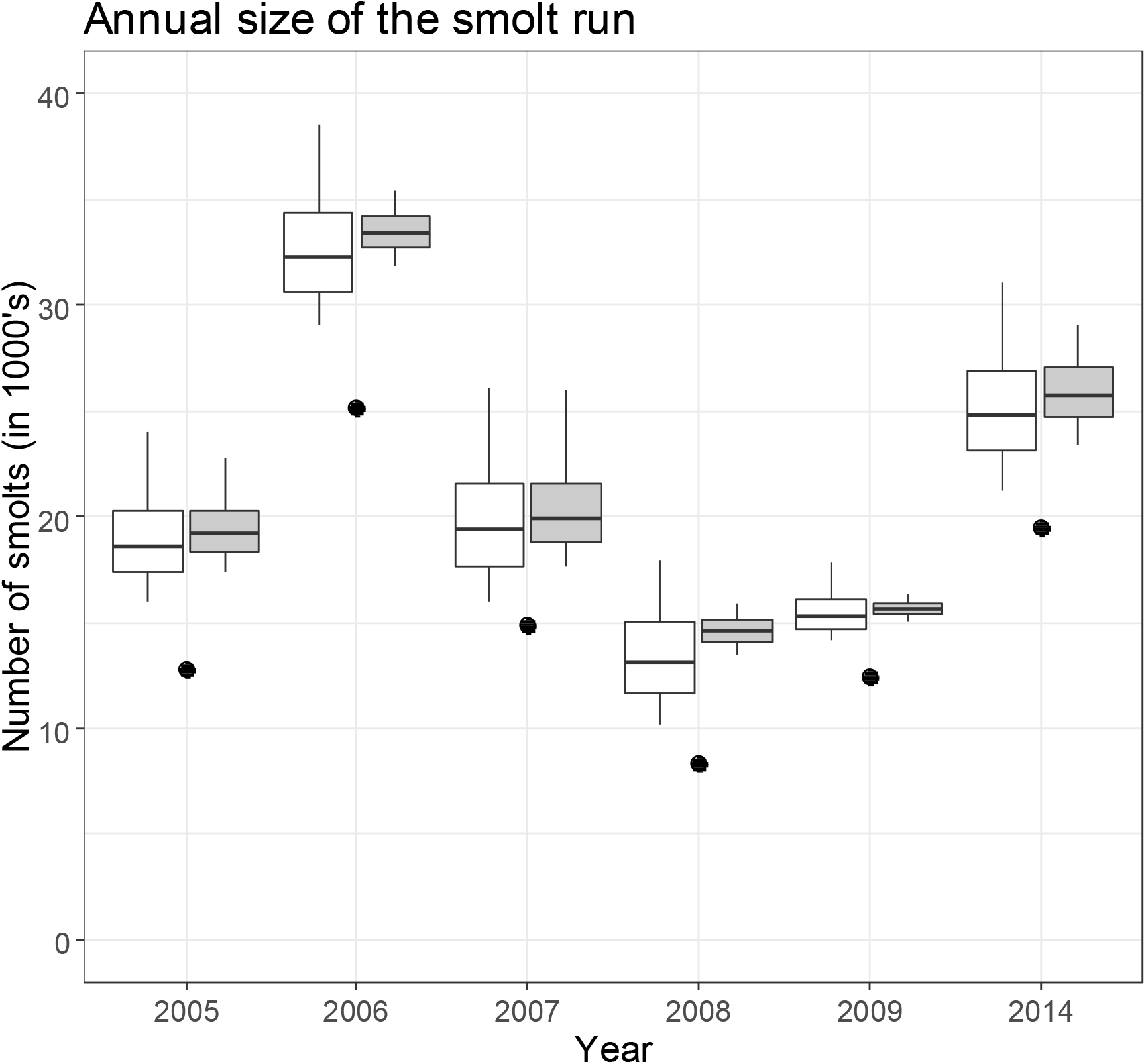
Posterior estimates of annual number of smolts passing the video site based on model run with expert elicited priors (grey boxplots) and priors from sensitivity analysis (white boxplots). Middle line is the median of the distribution; box covers 50% and whiskers 90% of the probability mass. Count data are illustrated with dots.

**Figure B5.**
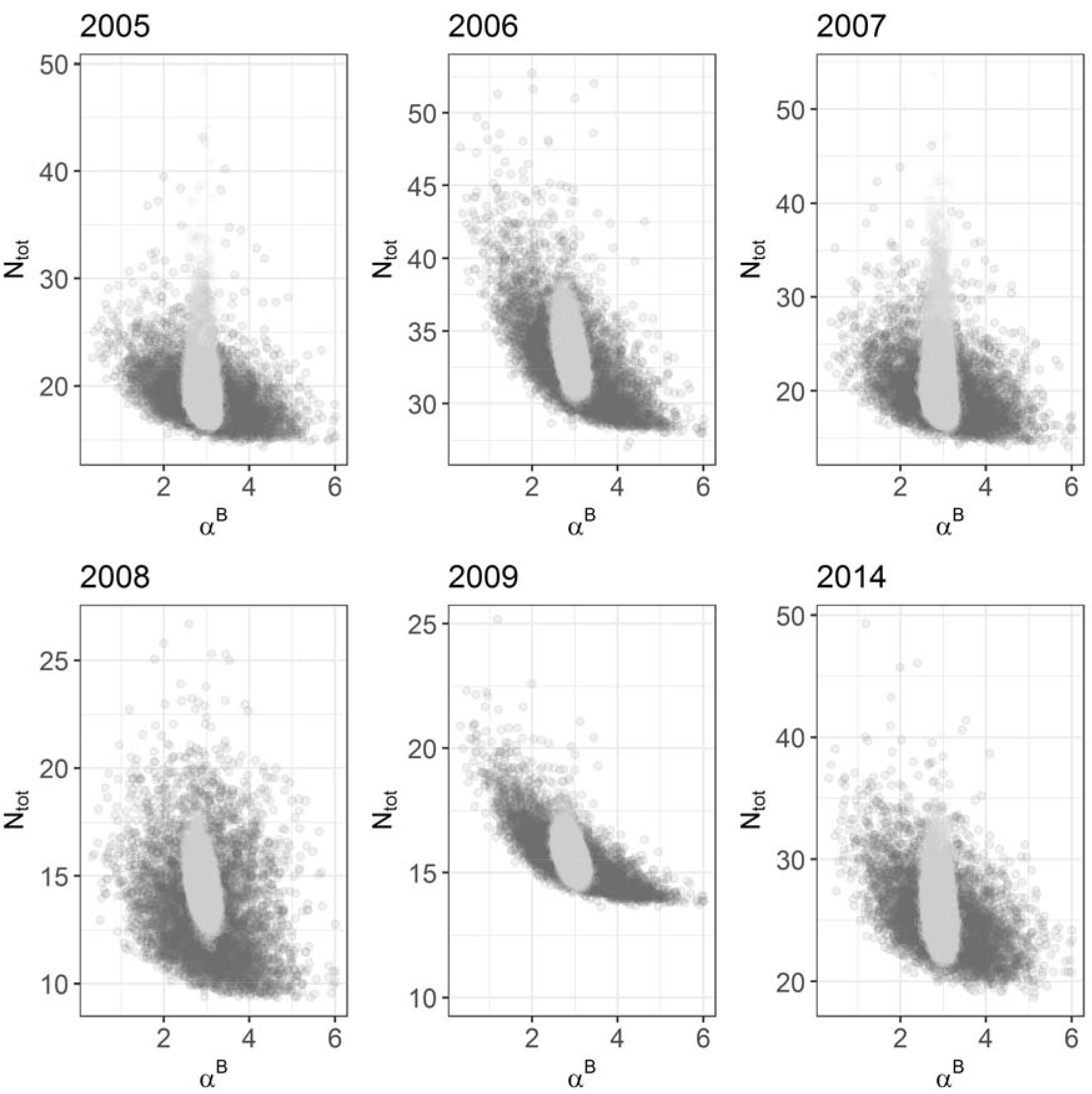
Scatter plots of sampled posterior values between annual total numer of smolts (in thousands) and parameter *α^B^* of the observation process. Dark grey dots illustrate the posterior samples from the sensitivity analysis, whereas light grey dots illustrate the posterior samples from the analysis with expert elicited priors.

**Figure B6.**
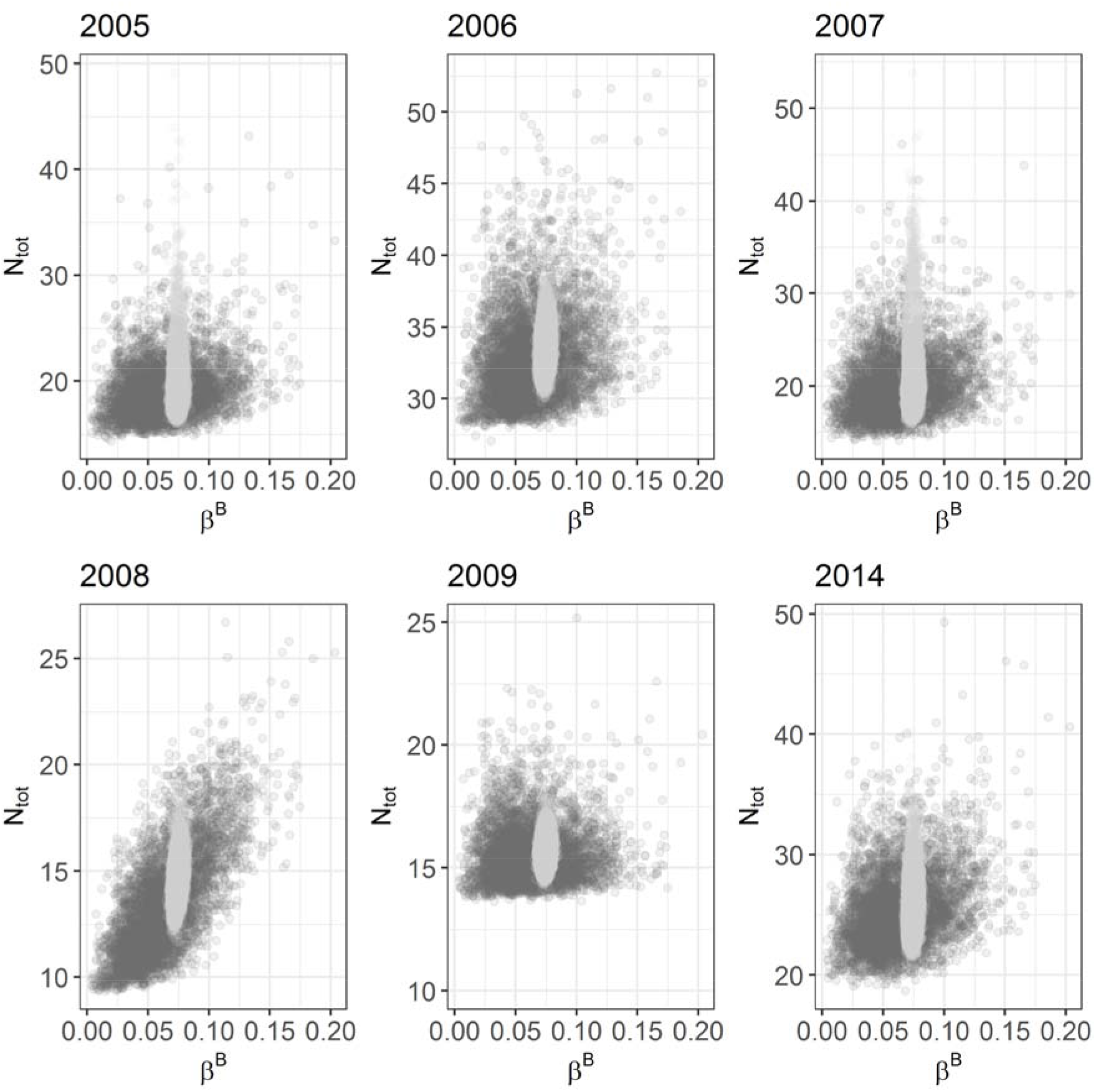
Scatter plots of sampled posterior values between annual total numer of smolts (in thousands) and parameter *β^B^* of the observation process. Dark grey dots illustrate the posterior samples from the sensitivity analysis, whereas light grey dots illustrate the posterior samples from the analysis with expert elicited priors.

**Figure B7.**
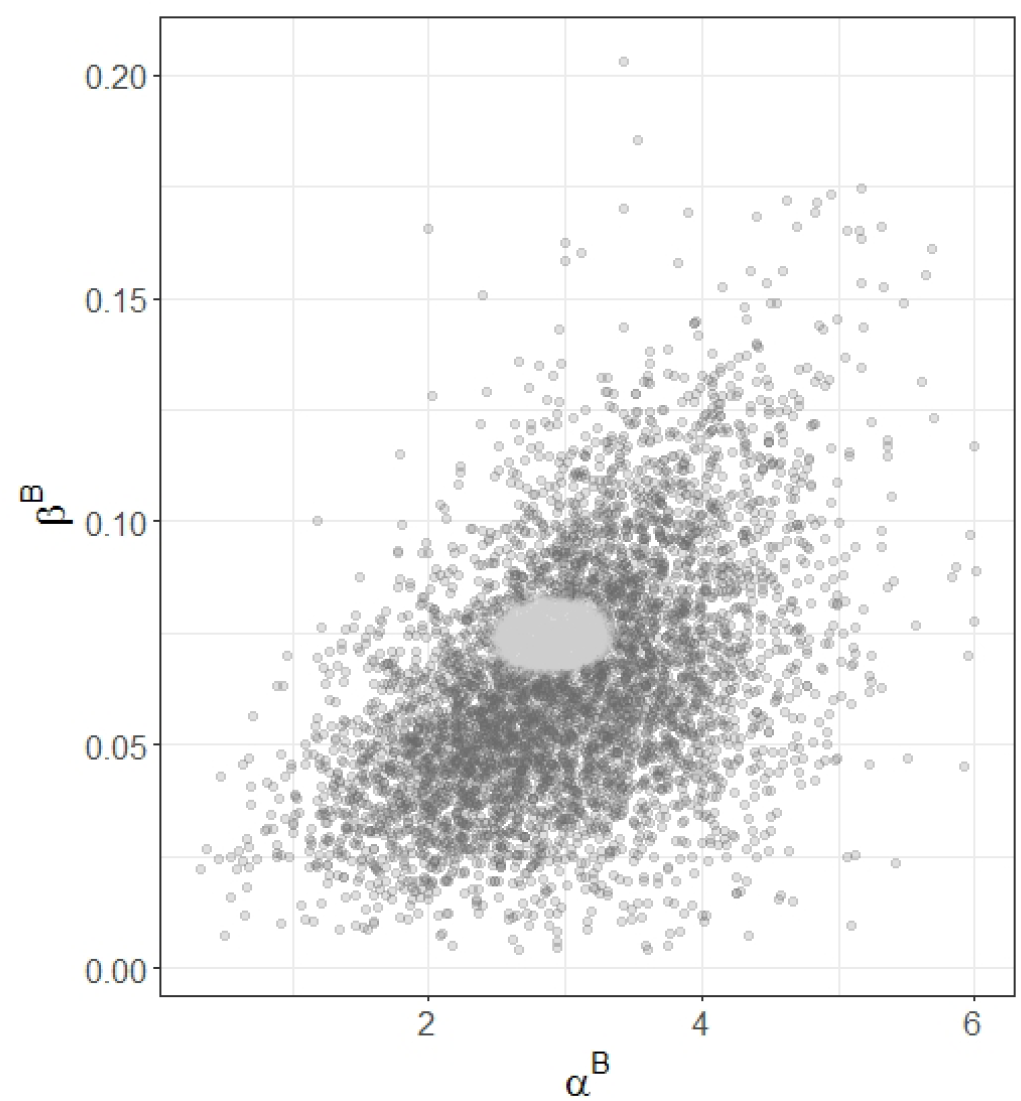
Scatter plots of sampled posterior of parameters *α^B^* and *β^B^* of the observation process. Dark grey dots illustrate the posterior samples from the sensitivity analysis, whereas light grey dots illustrate the posterior samples from the analysis with expert elicited priors.

## Appendix C: JAGS code for the arrival model

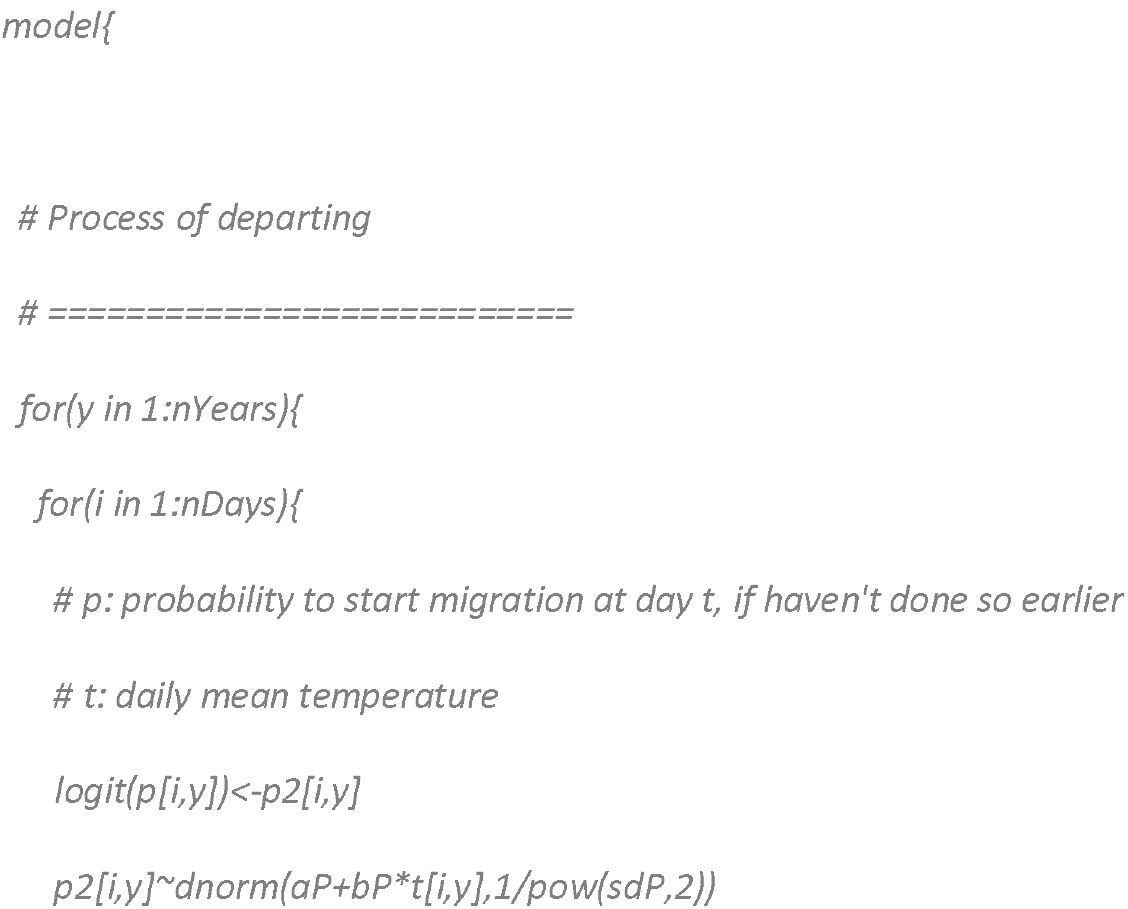
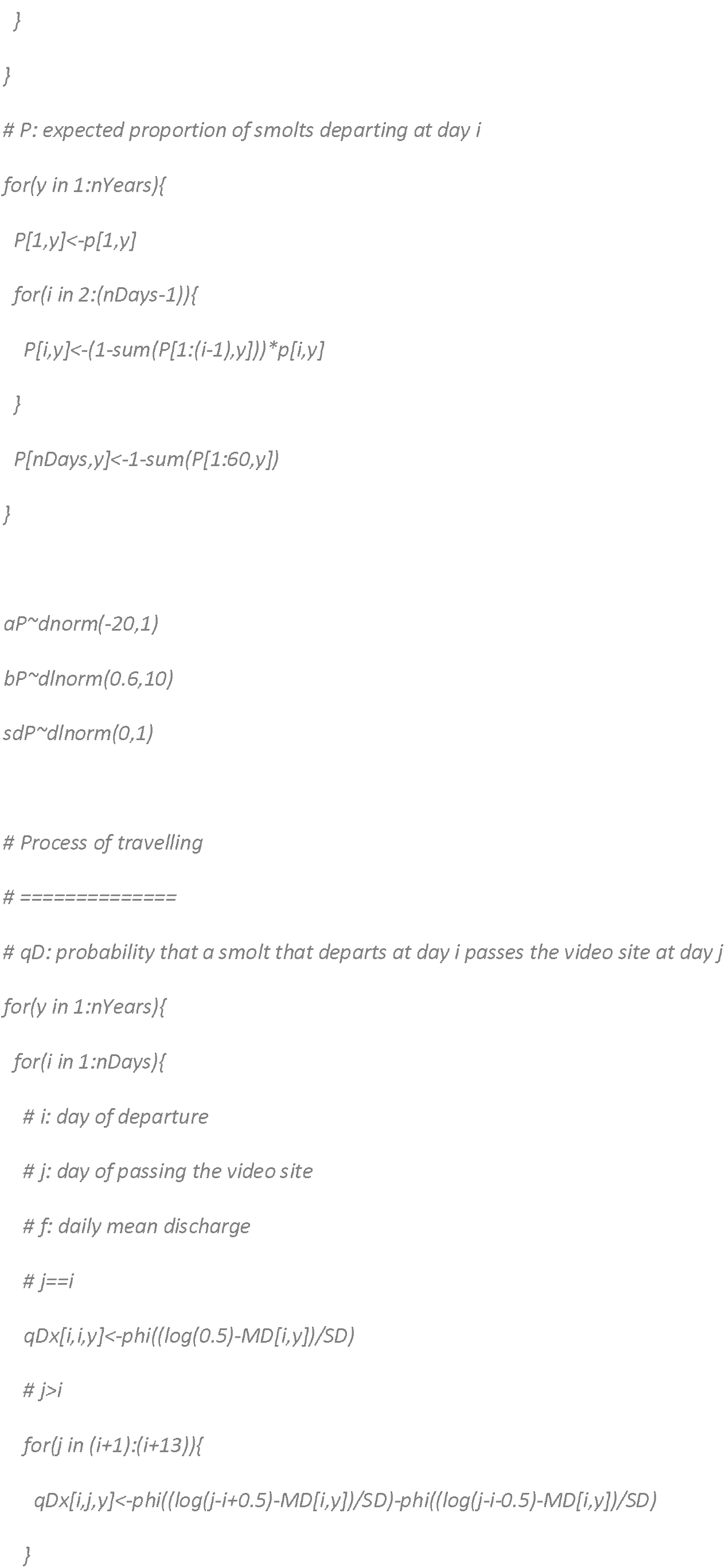
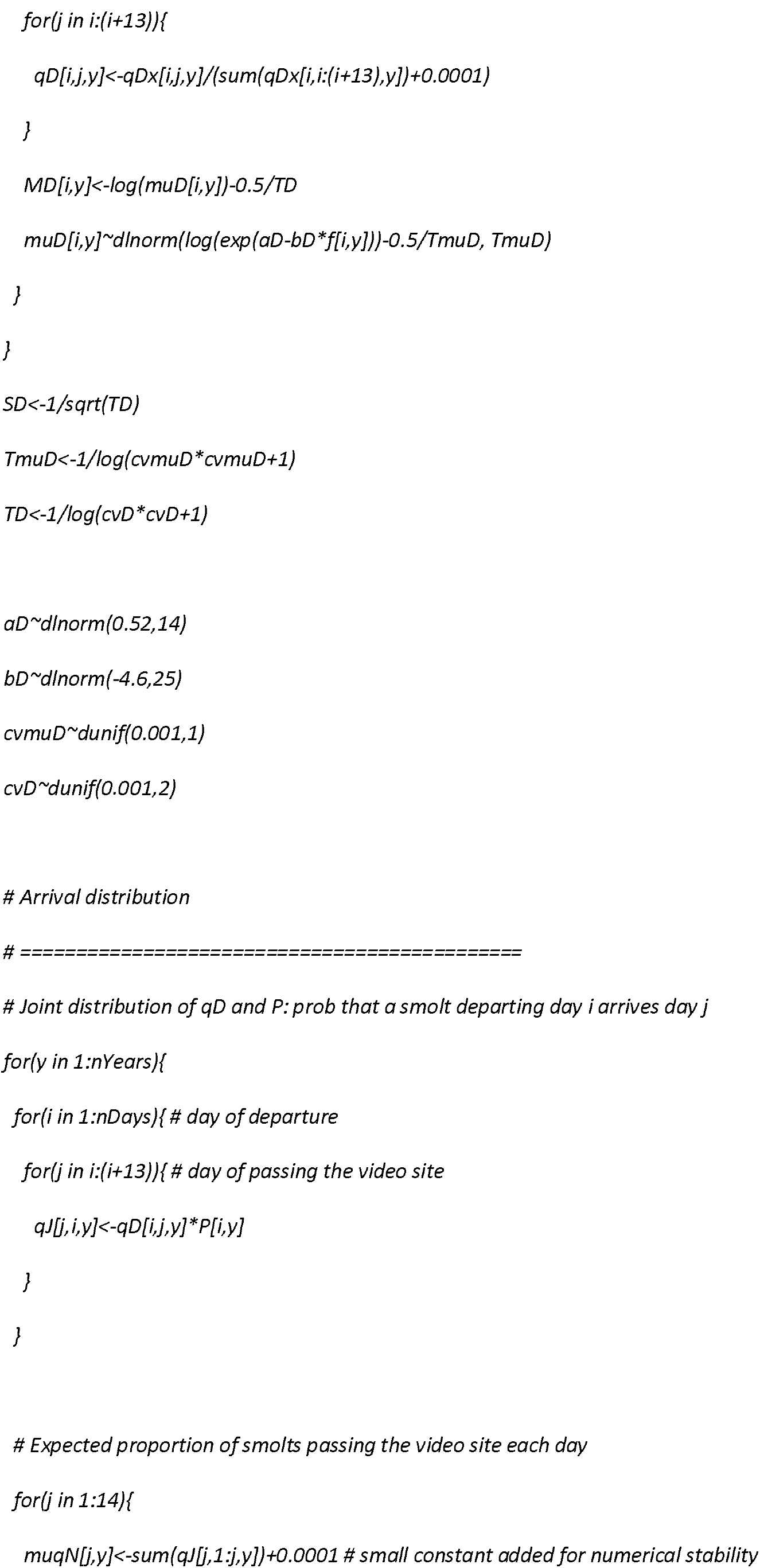
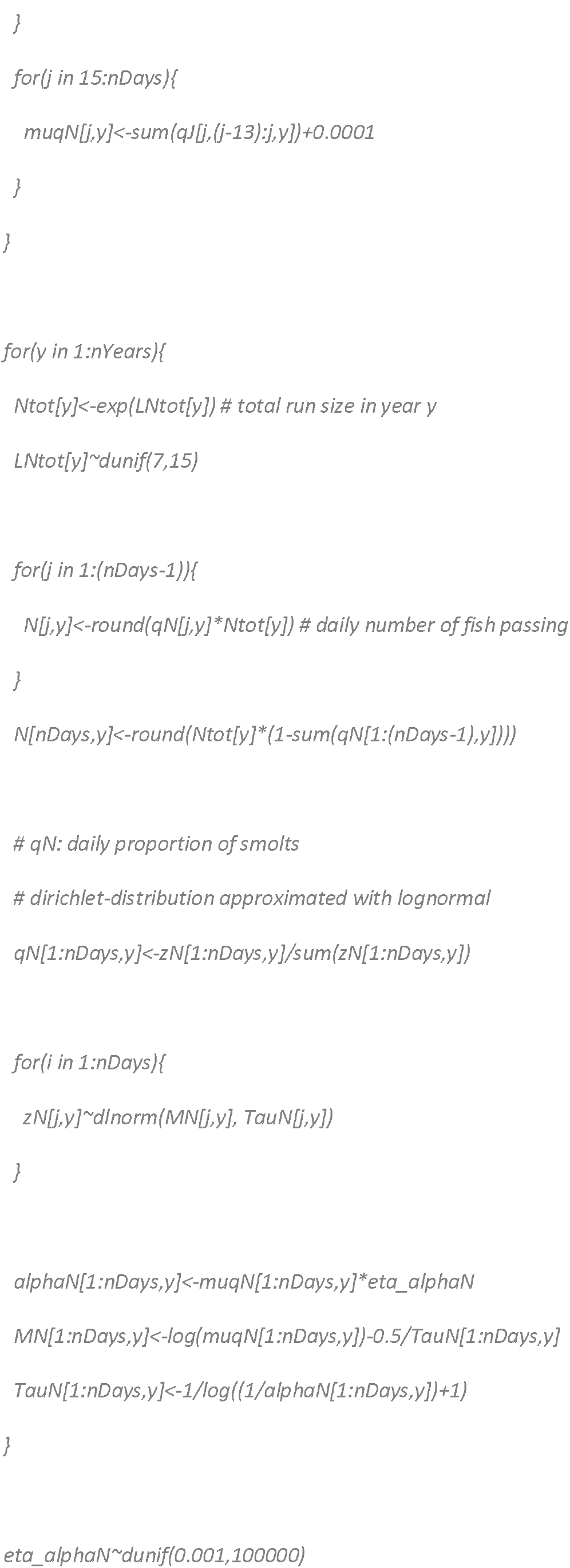
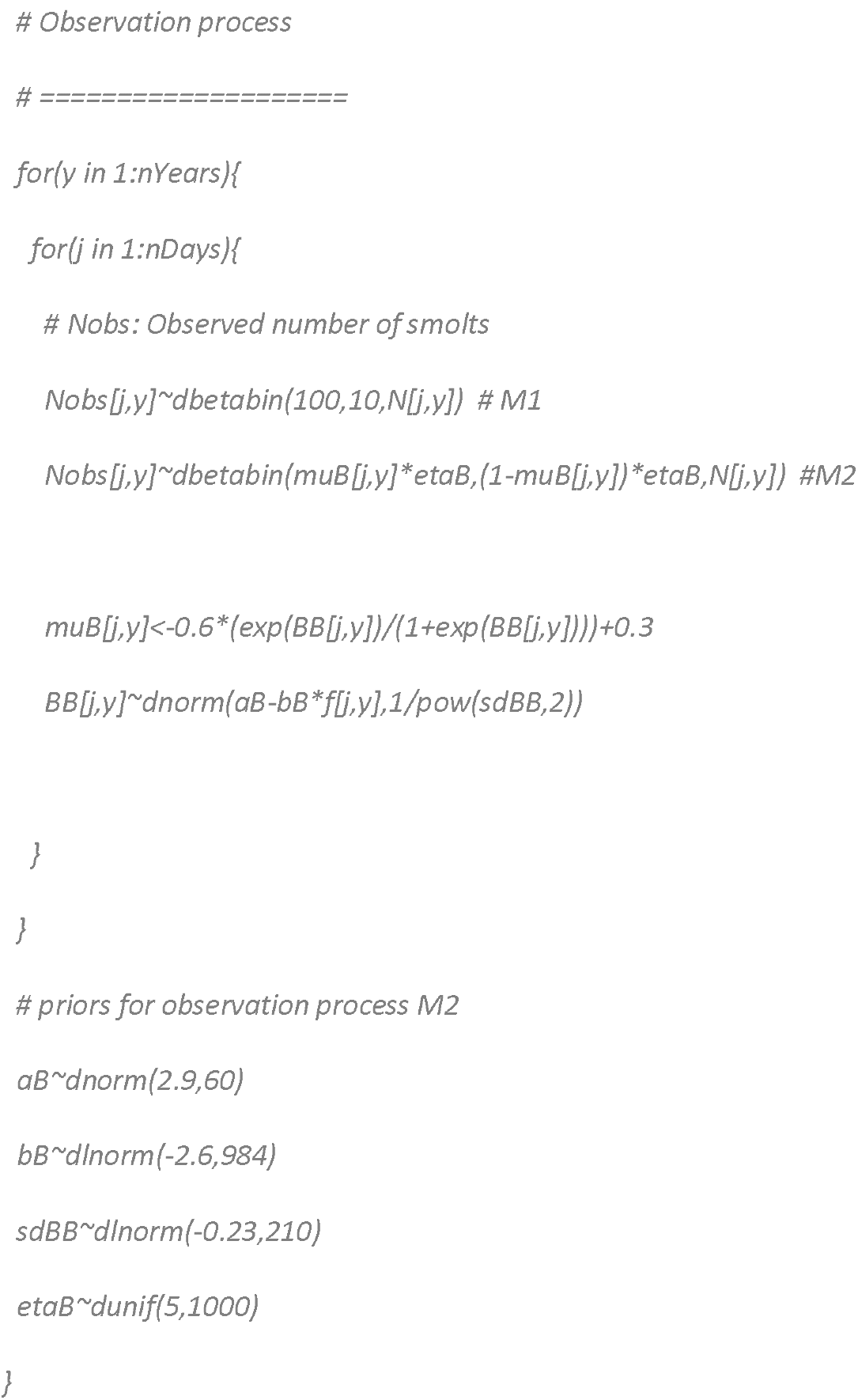

## References

Bakshtanskiy, E., Nesterov, V., and Neklyodov, M. 1988. Development of schooling behaviour in juvenile Atlantic salmon, Salmo salar, during seaward migration. J. Ichthyol. 28: 91–101.

Borgstrøm, R., Opdahl, J., Svenning, M.-A., Länsman, M., Orell, P., Niemelä, E., Erkinaro, J. & Dempson, J. B. 2010. Temporal changes in ascendance and is-season exploitation of Atlantic salmon, Salmo salar, inferred by a video camera array. Fish. Manage. Ecol. 17: 454–463.

Davidsen, J., Svenning, M. A., Orell, P., Yoccoz, N., Dempson, J. B., Niemelä, E., Klemetsen, A., Lamberg, A., and Erkinaro, J. 2005. Spatial and temporal migration of wild Atlantic salmon smolts determined from a video camera array in the sub-Arctic River Tana. Fish. Res. 74: 210–222.

Dempson, J. B, and Stansbury, D. E. 1991. Using partial counting fences and atwo-sample stratified design for mark–recapture estimation of an Atlantic salmonsmolt population. N. Am. J. Fish. Manage. 11: 27–37.

Denwood, M.J. 2016. runjags: An R Package Providing Interface Utilities, Model Templates, Parallel Computing Methods and Additional Distributions for MCMC Models in JAGS. J. Stat. Softw. 71(9): 1–25. doi:10.18637/jss.v071.i09.

Erkinaro, J., Julkunen, M. and Niemelä, E. 1998. Migration of juvenile Atlantic salmon Salmo salar in small tributaries of the subarctic River Teno, northern Finland. Aquaculture 168: 105–119.

Erkinaro, J., Czorlich, Y., Orell, P., Kuusela, J., Länsman, M., Falkegård, M., Pulkkinen, H., Primmer, C. & Niemelä, E. 2019. Life history variation across four decades in a diverse population complex of Atlantic salmon in a large subarctic river. Can. J. Fish. Aquat. Sci. 76: 42–55.

Gelman, A., and Rubin, D.B. 1992. Inference from iterative simulation using multiple sequences. Stat. Sci. 7: 457–511.

Hilborn, R., Bue, B.G., and Sharr, S. 1999. Estimating spawning escapements from periodic counts: a comparison of methods. Can. J. Fish. Aquat. Sci. 56(5): 888–896. doi:10.1139/f99-013.

Hansen, L. P., and Jonsson, B. 1985. Downstream migration of hatchery-reared smolts of Atlantic salmon (Salmo salar L.) in the River Imsa. Aquaculture, 45: 237–248.

Holmes, J.A, Cronkite, G.M.W., Enzenhofer, H. J., and Mulligan, T.J. 2006. Accuracy and precision of fish-count data from a “dual-frequency identification sonar” (DIDSON) imaging system. ICES J. Mar. Sci. 63: 543–555. doi:10.1016/j.icesjms.2005.08.015

Kuparinen, A., Mäntyniemi, S., Hutchings, J.A., and Kuikka, S. 2012. Increasing biological realism of fisheries stock assessment: towards hierarchical Bayesian methods. Environ. Rev. 20: 135–151. doi:10.1139/A2012-006.

Michielsens, C.G.J., McAllister, M.K., Kuikka, S., Mäntyniemi, S., Romakkaniemi, A., Pakarinen, T., Karlsson, L, and Uusitalo, L., 2008. Combining multiple Bayesian data analyses in a sequential framework for quantitative fisheries stock assessment. Can. J. Fish. Aquat. Sci. 65: 962–974. doi:10.1139/F08-015

Mäntyniemi, S., and Romakkaniemi, A. 2002. Bayesian mark-recapture estimation with an application to a salmonid smolt population. Can. J. Fish. Aquat. Sci. 59: 1748–1758. doi:10.1139/F02-146.

Mäntyniemi, S.H.P., Whitlock, R.E., Perälä, T.A., Blomstedt, P.A., Vanhatalo, J.P., Rincon, M.M., Kuparinen, A.K., Pulkkinen, H.P., and Kuikka, O.S. 2015. General state-space population dynamics model for Bayesian stock assessment. ICES J. Mar. Sci. 72: 2209–2222. doi:10.1093/icesjms/fsv117

Mäntyniemi, S., Kuikka, S., Rahikainen, M., Kell, L.T., Kaitala, V. 2009. The value of Information in fisheries management: North Sea herring as an example. ICES J. Mar. Science 66: 2278–2283.

Orell, P., Erkinaro, J., Svenning, M., Davidsen, J., and Niemelä, E. 2007. Synchrony in the downstream migration of smolts and upstream migration of adult Atlantic salmon in the sub-Arctic River Utsjoki. J. Fish. Biol. 71: 1735–1750.

Orell, P., Erkinaro, J., and Karppinen, P. 2011. Accuracy of snorkelling counts in assessing spawning stock of Atlantic salmon, Salmo salar, as verified by radio-tagged fish and underwater video monitoring. Fish. Manag. Ecol. 18: 392–399.

Otero, J., L’Abée-Lund, J.H., Castro-Santos, T., Leonardsson, K., Storvik, G.O., Jonsson, B., Dempson, B., Russell, I.C., Jensen, A.J., Baglinière, J.-L., Dionne, M., Armstrong, J.D., Romakkaniemi, A., Letcher, B.H., Kocik, J.F., Erkinaro, J., Poole, R., Rogan, G., Lundqvist, H., MacLean, J.C., Jokikokko, E., Arnekleiv, J.V., Kennedy, R.J., Niemelä, E., Caballero, P., Music, P.A., Antonsson, T., Gudjonsson, S., Veselov, A.E. Lamberg, A., Groom, S., Taylor, B.H., Taberner, M., Dillane, M., Arnason, F., Horton, G., Hvidsten, N.A., Jonsson, I.R., Jonsson, N., McKelvey, S., Næsje, T., Skaala, ø., Smith, G.W., Sægrov, H., Stenseth, N.C. and Vøllestad, L.A. 2014. Basin-scale phenology and effects of climate variability on global timing of initial seaward migration of Atlantic salmon (Salmo salar). Glob. Change Biol. 20: 61–75.

Plummer, M. 2003. JAGS: A program for analysis of Bayesian graphical models using Gibbs sampling. In proceedings of the 3^rd^ International Workshop on Distributed Statistical Computing (DSC 2003), 20-22 March, Vienna, Austria. ISSN 1609-395X.

Romakkaniemi, A., Lilja, J., Nykänen, M., Marjomäki, T.J. and Jurvelius, J. 2000. Spawning run of Atlantic Salmon (Salmo salar) in the River Tornionjoki monitored by horizontal split-beam echosounding. Aquat. Living Resour. 13: 349–354.

Schwarz, C., and Dempson, J. 1994. Mark-recapture estimation of a salmon smolt population. Biometrics. 50: 98–108.

Sethi, A.S., and Bradley, C. 2016. Statistical arrival models to estimate missed passage counts at fish weirs. Can. J. Fish. Aquat. Sci. 73: 1251–1260. doi:10.1139/cjfas-2015-0318.

Spiegelhalter, D.J., Best, N.G., Carlin, B.P., and van der Linde, A. 2002. Bayesian measures of model complexity and fit (with discussion). J. R. Stat. Soc. Ser. B. 64: 583–639.

Su, Z., Adkison, M.D., and Van Alen, B.W. 2001. A hierarchical Bayesian model for estimating historical salmon escapement and escapement timing. Can. J. Fish. Aquat. Sci. 58: 1648–1662. doi:10.1139/f01-099.

Trépanier, S., Rodríquez, M. A. and Magnan, P. 1996. Spawning migrations in landlocked Atlantic salmon: time series modelling of river discharge and water temperature effects. J. Fish Biol. 48: 925–936.

Laura Uusitalo, Sakari Kuikka, Atso Romakkaniemi; Estimation of Atlantic salmon smolt carrying capacity of rivers using expert knowledge, ICES Journal of Marine Science, Volume 62, Issue 4, 1 January 2005, Pages 708–722, https://doi.org/10.1016/j.icesjms.2005.02.005

Vähä, J.P., Erkinaro, J., Falkegård, M., Orell, P. & Niemelä, E. 2017. Genetic stock identification of Atlantic salmon and its evaluation in a large population complex. Can. J. Fish. Aquat. Sci. 74: 327–338.

